# A human iPS cell line for ready-to-use human iAstrocytes that support human neurons

**DOI:** 10.64898/2026.03.02.709009

**Authors:** Larissa Breuer, Hanna Dubrovska, Jeremy Krohn, John Carl Begley, Hana Toda Sheldon, Katarzyna A. Ludwik, Harald Stachelscheid, Camin Dean

## Abstract

Human iPSC-derived neuronal networks are increasingly being employed in basic and applied research to enhance translation. Astrocytes are essential for neuronal network function, but are often not included, or replaced with mouse astrocytes, which compromises translation. Current protocols produce hiPSC-derived astrocytes by stepwise differentiation using small molecules and cytokines, or by forward programming by inducing transcription factors introduced by lentiviral transduction. Here we created a stable, inducible hiPSC line capable of producing iAstrocytes by introducing the transcription factors NFIB and SOX9 into the AAVS1 locus of the BIHi005-A hiPSC line. iAstrocytes induced from this line upregulated astrocytic genes over four weeks in culture, expressed GFAP and S100B and exhibited spontaneous calcium waves and responses to ATP and CPA. In co-cultures, iAstrocytes supported the growth and function of mature iNeuron networks. Pre- and post-synaptic markers and synchronous neuronal activity measured by high-density multi-electrode array recordings and neuronal calcium imaging, appeared by four weeks. The use of iAstrocytes will help to standardize the use of human astrocytes to support human neural networks and enhance translation.

## Introduction

Human induced pluripotent stem cell (hiPSC)-derived neuronal networks are increasingly being used in both basic and applied research to address the translational gap between preclinical findings in rodent and the treatment of human neurological diseases (Marshall et al., 2023). However, hiPSC-based models are often limited in complexity due to a lack of functional human astrocytes, which are essential for synapse formation and function (Chung et al., 2015) and the development of mature functional networks (Engle et al., 2018).

The most common hiPSC-based culture systems use differentiation by small molecules and cytokines or forward programming by induction of transcription factors. Small molecule approaches usually take longer. For example, dual SMAD inhibition by small molecules induces cortical glutamatergic neurons via neural progenitor cells (NPCs) in approximately 6 weeks (Chambers et al., 2009). Astrocytes are not present initially, but NPCs can develop into astrocytes between 5 and 8 weeks, especially if serum or astrocyte-promoting factors are added to the media. Doxycycline-induced expression of the transcription factor NGN2 can more rapidly and efficiently turn hiPSCs into neurons (iNeurons), in less than two weeks (Zhang et al., 2013). To support iNeuron growth and maturation, mouse astrocytes were added (Zhang et al., 2013). Subsequent studies reported that co-culturing iNeurons with rodent astrocytes (Meijer et al., 2019; Rhee et al., 2019) or with rodent astrocyte conditioned media (Shan et al., 2024) enhances iNeuron network function. One of these studies reported that rodent astrocytes are superior to human astrocytes in an autaptic model (Rhee et al., 2019), initially discouraging efforts towards a fully human system.

However, mouse astrocyte support of human neurons could still severely compromise translation, given the amount of crosstalk between astrocytes and neurons, especially in disease states (Linnerbauer et al., 2020). For example, mouse and human astrocytes respond differently to inflammation (Tarassishin et al., 2014), a common feature of many brain disorders. This likely stems from species-specific differences in gene expression. Glial cells are the least conserved brain cells between human and mouse (Pembroke et al., 2021) with microglia being the most divergent, followed by astrocytes, which are only 50-60% similar between human and mouse (Zhang et al., 2016). Given the central role astrocytes play in both healthy and diseased brain states (Lee et al., 2022; Oksanen et al., 2020; Phatnani and Maniatis, 2015), incorporating human astrocytes into human-based neuronal model systems is essential for translation. Our goal, therefore, was to create a gene-edited human iPSC line to generate human astrocytes that support human neurons in culture.

The transcription factors SOX9, NFIA, and NFIB, rapidly convert human iPSCs or fibroblasts to an astrocytic fate. These factors have been used by lentiviral transduction alone or in combination by different studies (Caiazzo et al., 2015; Canals et al., 2018; Li et al., 2018; Neyrinck et al., 2021; Yeon et al., 2021; Baranes et al., 2023). SOX9 initiates gliogenesis and induces expression of NFIA and NFIB, which promote astrocyte differentiation (Deneen et al., 2006; Kang et al., 2012). Since lentiviral transduction of inducible SOX9/NFIB (Canals et al., 2018), and SOX9/NFIA (Li et al., 2018) both resulted in robust, fast, efficient induction of human astrocytes, we initially created hiPSC lines by introducing these transcription factor combinations. We found that our inducible SOX9/NFIB hiPSC line produced functional and mature human astrocytes (iAstrocytes), validated by gene expression, astrocytic marker protein expression, and calcium signaling. We compared NGN2-induced iNeuron network function without astrocyte support, with mouse astrocytes, or with our iAstrocytes in immunostains of synaptic markers, and using multi-electrode arrays and calcium imaging. We found that iAstrocytes support iNeuron growth, synapse formation, and network function and maturation. This gene-edited hiPSC line therefore provides iAstrocytes that support human neurons and can be used to advance translation of basic research to new clinical treatments of brain diseases and disorders.

## Results

### Astrocyte markers are upregulated in iAstrocytes

We first generated a gene-edited hiPSC line by introducing a construct encoding inducible NFIB/SOX9 (Fig. 1A) into the AAVS1 locus of the BIHi005-A hiPSC line. To test if the induced gene-edited puromycin-selected polyclonal hiPSCs differentiated into astrocytes, we examined a time course of gene expression in samples collected the day before induction, and on days 3, 6, 13, 20 and 27 after induction with doxycycline on day 1, where day 21 is assumed to be “mature” astrocytes (Canals et al., 2018). The most down-regulated genes were related to pluripotency and include NANOG and POU5F1 (OCT4) (Fig. 1B), indicating that the polyclonal NFIB/SOX9 hiPSCs left the stem cell state progressively during the time course. The top 25 up-regulated genes are all found in astrocytes (Stelzer et al., 2016), except for GDF6 and ACKR1, which are mainly present in oligodendrocyte precursor cells and endothelial cells, respectively. The expression of the induced transcription factors NFIB and SOX9 increased by DoI 3 and remained elevated throughout later time points (Fig. 1C), indicating that induction was successful and maintained over time. In an examination of astrocytic markers, S100B, GFAP, AQP3, PAX6, FABP7, and SLC1A2 (EAAT2) were upregulated, but there was no change in ALDH1L1 or SLC1A3, and NFIA and AQP4 were downregulated (Fig. 1C). S100B was the most upregulated astrocyte-specific marker, suggesting that mature astrocytes were induced (Raponi et al., 2007). There was an overall general drop in gene expression on Day 6 of the time course, which may have been caused by splitting and treating the cells with ROCK inhibitor, such that differentiation underwent a brief reset from which the cells had to recover.

**Figure 1.**
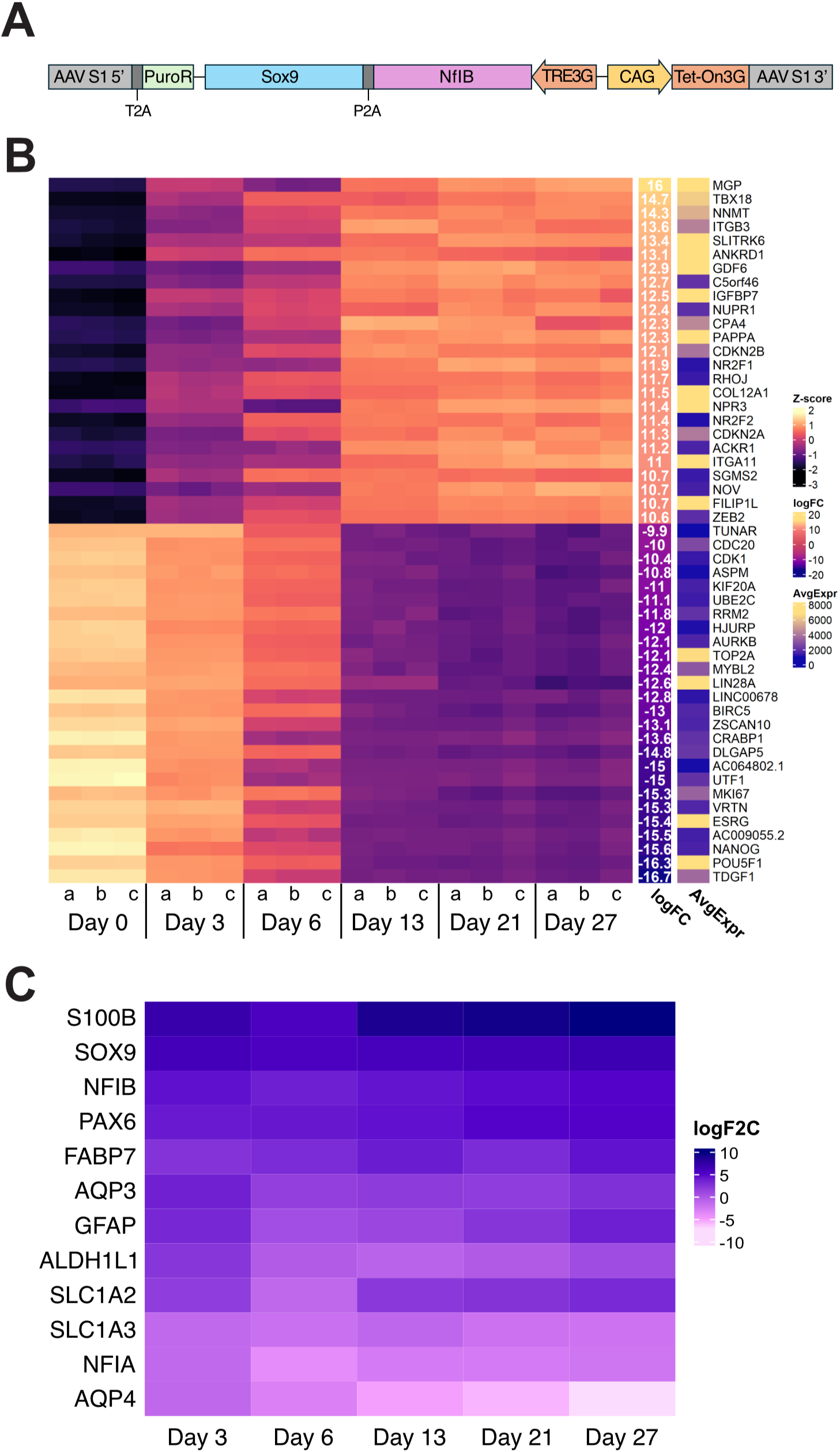
NFIB+SOX9 induced cells increase astrocytic markers and decrease pluripotency markers. **A)** The construct used to create the NFIB+SOX9 inducible hiPSC line contained flanking homology arms for the AAVS1 locus, a Puromycin resistance gene, Tet-On3G driven by the CAG promoter, and NFIB and SOX9 separated by a P2A site and driven by the TRE3G promoter. **B)** Top 25 up and downregulated differentially expressed genes (Log fold change) with an average expression of ≥ 1000 normalized counts in three technical replicates (a, b, and c) throughout differentiation of the NFIB/SOX9 polyclonal hiPSC line at day of induction 0, 3, 6, 13, 20, and 27. **C)** NFIB and SOX9 expression is increased by doxycycline induction, and astrocytic gene expression is upregulated over time in the induced NFIB/SOX9 polyclonal hiPSC line. Log2-fold change of gene expression compared to Day 0 before induction is shown.

### iAstrocytes support iNeurons and synapses in co-cultured iNeurons

iAstrocytes generated from the polyclonal gene-edited hiPSCs were then tested for their ability to support iNeurons in culture. The NGN-2 inducible hiPSC line BIHi005-A-24 was pre-differentiated for seven days to neural stem cells (NSCs), which were plated at a density of 100,000 cells per well in poly-D-lysine/Biolaminin-coated 24-well plates in Modified Neural Expansion Medium. The day after plating the medium was changed to iNeuron Medium, with daily full medium changes thereafter for the next six days. On day 7, iAstrocytes, pre-differentiated for seven days, were added at a density of 50,000 cells per well to iNeurons, and the media changed to Co-Culture Medium. After a full medium change the day after addition of iAstrocytes, half of the medium was changed every 2-3 days, always containing fresh doxycycline, for maintenance of cultures. Cultures were treated with AraC for 24 hours two to four days after adding iAstrocytes to limit astrocytic growth. We found that adding iAstrocytes that were doxycycline-induced and pre-differentiated for seven days to iNeurons induced from NSCs for seven days, was optimal for survival of both cell types. iNeurons preferentially grew on differentiated iAstrocytes (Fig. 2) compared to areas of undifferentiated cells, indicating that iAstrocytes support iNeurons in co-culture.

**Figure 2.**
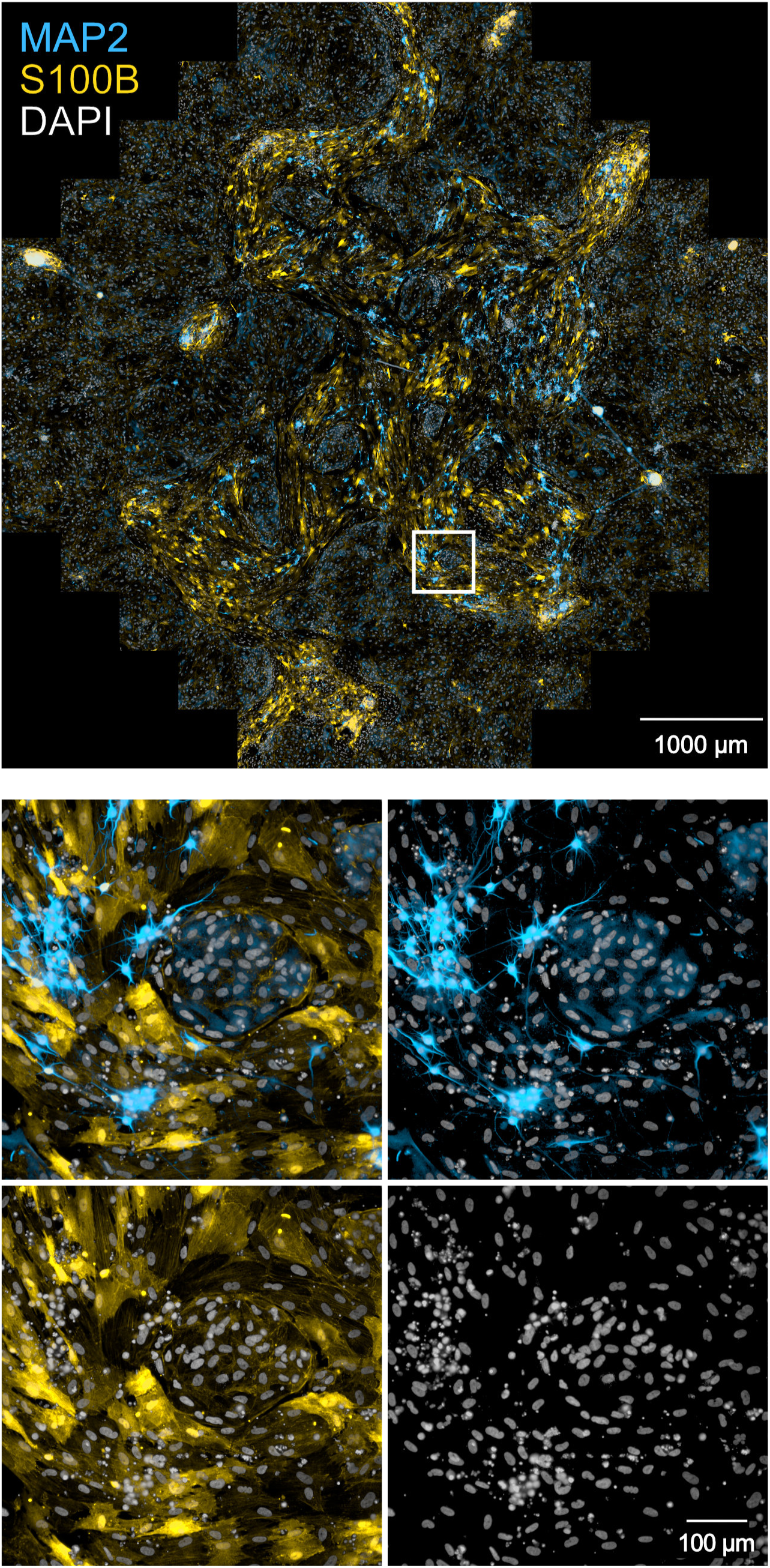
NFIB+SOX9 polyclonal iAstrocytes support iNeurons in co-culture. DIV21 co-cultured iNeurons and iAstrocytes (cultured separately for 7 days, then together for 14 days) immunostained for MAP2 (cyan), S100B (yellow), and DAPI (white). Whole well view shows that MAP2-positive iNeurons preferentially grow on regions of iAstrocytes identified by S100-positive signal. Scale bar = 1 mm. Lower panels are a zoom of the indicated region (white square in top panel) showing a region where iAstrocytes (yellow) support iNeurons (cyan), compared to surrounding areas with only DAPI-positive cells. Scale bar =100 µm.

We next compared synapse markers in cultures of iNeurons alone, iNeurons co-cultured with mouse astrocytes (mAstrocytes), or iNeurons co-cultured with iAstrocytes. At DIV28, cultures showed prominent synapses stained by synaptophysin (Fig. 3A). Quantification of synapses per length dendrite along MAP2-positive processes showed no significant difference in synapse number between iNeurons co-cultured with iAstrocytes (0.23 ± 0.05 synapses/µm, mean ± SD) and iNeurons co-cultured with mouse astrocytes (0.24 ± 0.06 synapses/µm, mean ± SD; p = 0.52). However, both conditions with astrocytes had significantly more synapses than iNeurons alone (0.19 synapses/µm ± 0.04 synapses/µm; p < 0.0001; two-way ANOVA, Tukey’s test, n = 50 dendrites from two biological replicates). The presynaptic active zone marker protein Bassoon, and the post-synaptic marker Homer also co-localized at synapses at DIV28 (Fig. 3B), indicating mature pre- and post-synaptic specializations.

**Figure 3.**
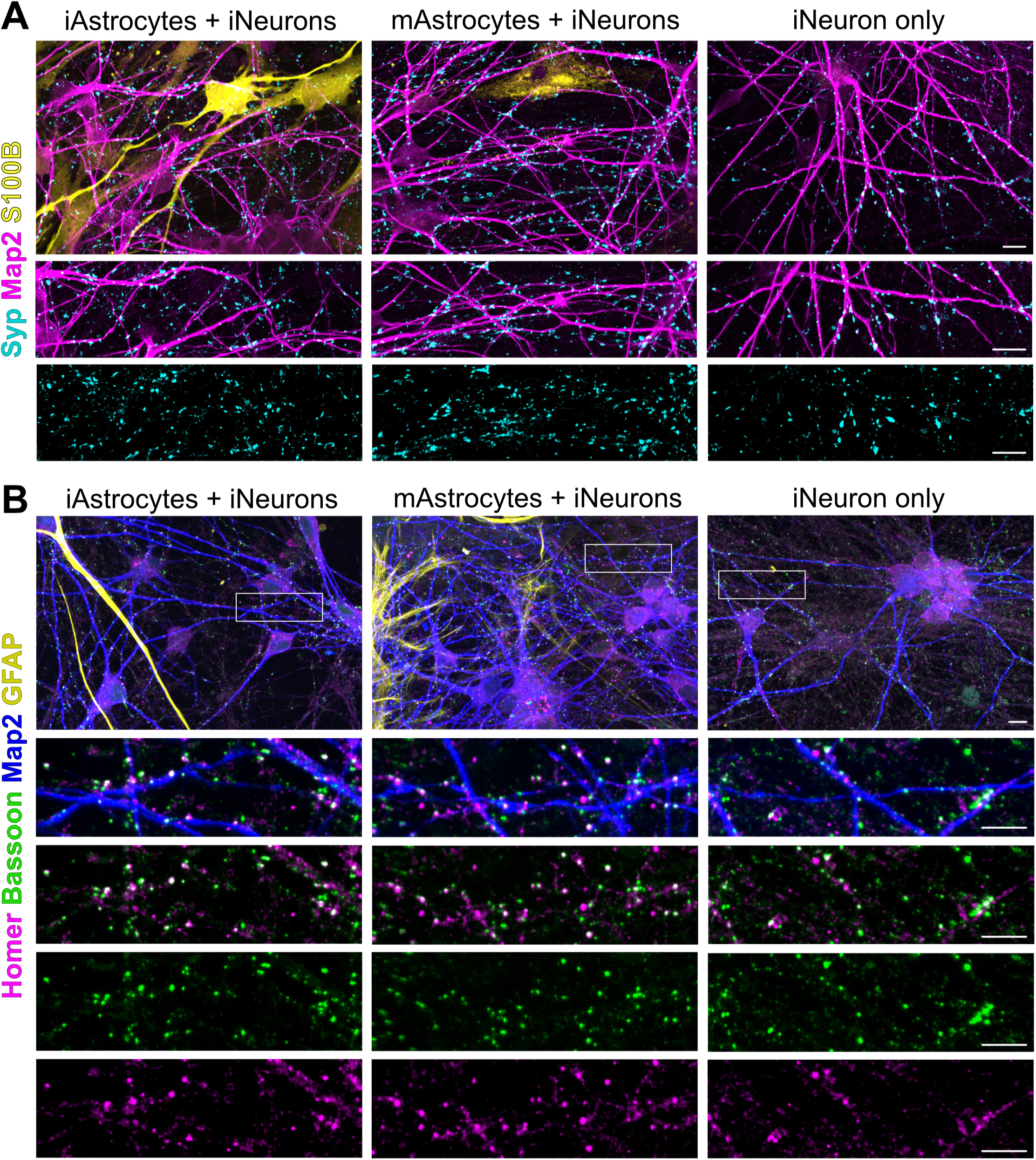
iAstrocytes support synapse formation in co-culture. **A)** DIV28 iAstrocytes + iNeurons, mouse astrocytes (mAstrocytes) + iNeurons, and iNeurons only without astrocytes, immunostained for S100B (yellow), synaptophysin (Syp; cyan) and MAP2 (magenta). Middle panel shows a zoom of Synaptophysin and MAP2 and bottom panel shows Synaptophysin alone. Scale bars = 10 µm. **B)** DIV28 iAstrocytes + iNeurons, mouse astrocytes (mAstrocytes) + iNeurons, and iNeurons only without astrocytes, immunostained for GFAP (yellow), Homer (magenta), Bassoon (green), and Map2 (blue). Bottom panels show a zoom of the indicated regions with Homer, Bassoon, and MAP2, Homer and Bassoon, Bassoon alone, and Homer alone. Scale bars = 10 µm.

### iAstrocytes from monoclonal gene-edited hiPSC lines support iNeurons

Given the promising results above, we proceeded to isolate monoclonal hiPSC lines from the polyclonal hIPSCs. Three clones, named BIHi005-A-1C (homozygous), BIHi005-A-1D (heterozygous), and BIHi005-A-1E (homozygous), registered on hPSCreg were chosen based on genotyping results and superior growth in culture after editing. We then tested if the chosen clones differentiated and matured as expected and if they supported iNeurons. iAstrocyte clones C, D, E, or mouse astrocytes (mAstrocytes) were co-cultured for 20 days with iNeurons, fixed on DIV27 of iNeuron / DIV29 of iAstrocyte induction, and immunostained with the neuronal marker MAP2, astrocytic markers GFAP and S100B, and DAPI. Although iAstrocytes did not grow to a confluent layer like mouse astrocytes did, in areas where iAstrocytes were present, we observed iNeurons thriving (Fig. 4A). iAstrocyte clone E showed good support of iNeurons, clone D showed moderate support, and clone C did not appear to support iNeurons. Quantification supported this observation. The proportion of MAP2-positive iNeurons relative to the overall detected cells (DAPI-positive) growing on clone E iAstrocytes reached 36.2% and 34.5%, for each of two repetitions, like iNeurons growing on mouse astrocytes (34.4% and 40.2%). Wells containing clone D iAstrocytes had slightly fewer iNeurons (26.7% and 19.1%), while wells containing clone C iAstrocytes only had 2.1% and 1.6% iNeurons (Fig. 4B).

**Figure 4.**
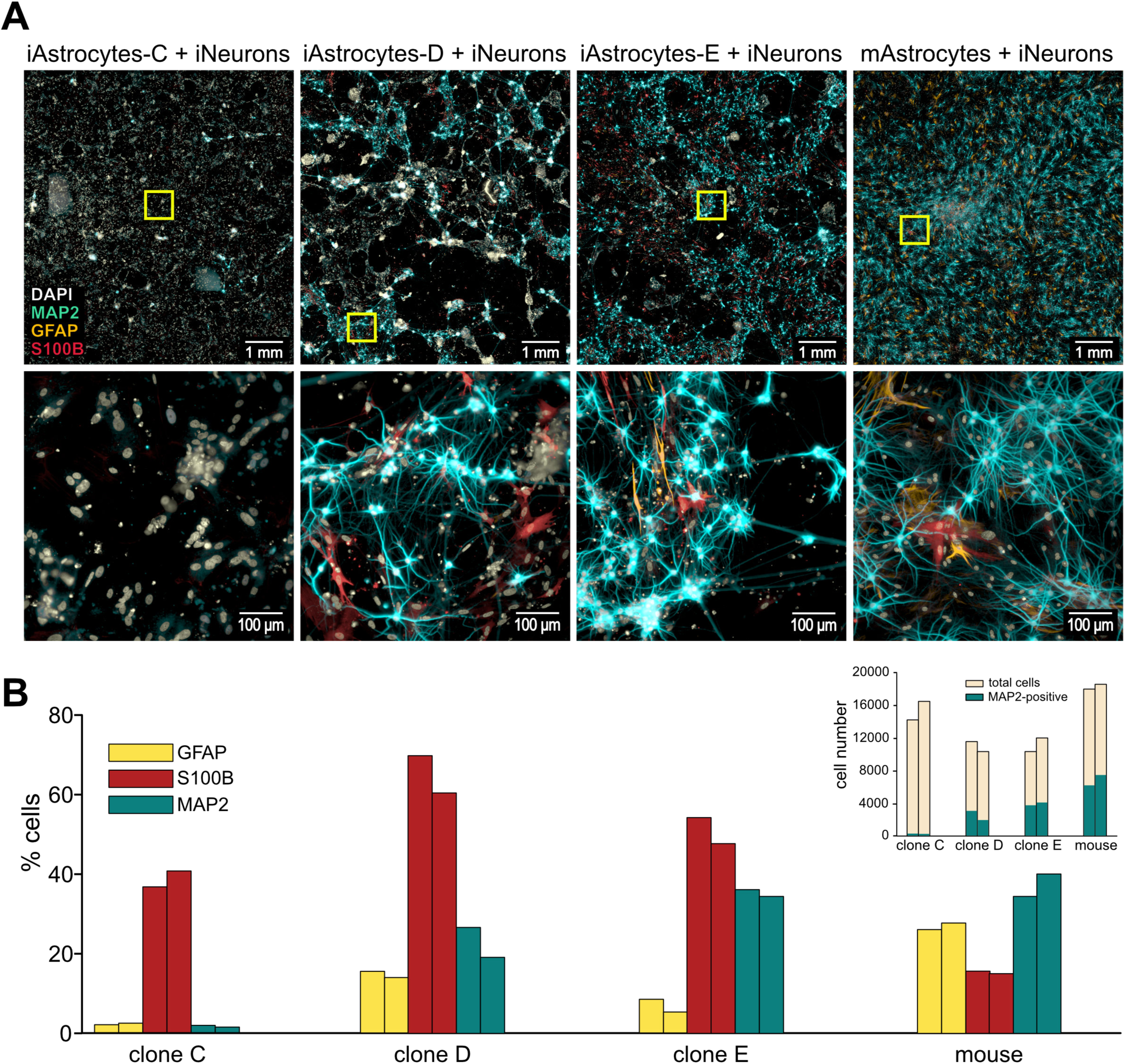
iAstrocyte from clones D and E support iNeurons in co-culture, but clone C does not. **A)** Whole-well view of one well of a 24-well plate of NFIB+SOX9 iAstrocyte clones C, D, E, or mouse astrocytes (mAstrocytes) co-cultured together for 20 days with iNeurons, fixed on DIV27 of iNeuron / DIV29 of iAstrocyte induction, and immunostained with MAP2 (cyan), GFAP (yellow), S100B (red), and DAPI (white). 10x zooms (lower panels) to fields of view indicated with yellow squares in top panels. **B)** Proportion of GFAP, S100B, and MAP2 positive cells relative to total DAPI-positive cells. The two bars per clone indicate technical replicates. Cells positive for one astrocytic marker could also be positive for the other marker, thus, assignment to one proportion of cells did not exclude assignment to the other proportion. Inset shows total cell count of DAPI-positive (white) and MAP2-positive cells per replicate and clone.

Quantitation of GFAP and S100B positive cells (by detection of GFAP or S100B signal surrounding DAPI-positive nuclei) revealed that the number of GFAP expressing cells was lower in iAstrocytes compared to mouse astrocytes while the number of S100B expressing cells was more than twice as high in iAstrocytes compared to mouse astrocytes (Fig. 4B). Most cells were either GFAP or S100B positive, but some cells expressed both markers (Fig. 4B). Mouse astrocyte cultures had more cells overall (Fig. 4B, inset), as did iAstrocyte clone C, possibly due to insufficient differentiation and proliferation of neural progenitor cells. Clone C was therefore omitted from further testing.

### iAstrocytes have spontaneous calcium events and respond to ATP and CPA

Astrocytes signal via calcium waves, and react to external stimuli such as adenosine triphosphate (ATP), which triggers calcium waves by activating purinergic receptors (Neary et al., 1988), and cyclopiazonic acid (CPA), which reduces calcium signaling in astrocytes by inhibiting the SERCA Ca2+ ATPase pump (Parri and Crunelli, 2003). We therefore tested iAstrocyte function by calcium imaging of spontaneous events and responses to ATP and CPA. iAstrocyte D and E clones transduced with AAV5 GFAP promoter-driven membrane-targeted GCaMP6f, exhibited spontaneous calcium events and responses to ATP (Fig. 5A, B). This confirmed that GFAP is expressed in a small subset of astrocytes, and these astrocytes are functional. To visualize all astrocytes, iAstrocyte clone E cultures - which supported iNeurons best - were treated with bath-applied Cal520 calcium dye on the day of imaging. iAstrocytes (clone E) exhibited spontaneous calcium waves (Fig. 5C). They also responded to ATP with an increased calcium response (Fig. 5D). AQuA software (Wang et al., 2019) was used to detect and quantify events, where a significant increase in calcium event area was found in response to ATP treatment (Fig. 5D). In addition, iAstrocytes responded to CPA with a decrease in calcium signaling, as expected, resulting in a significant decrease in frequency of calcium events (Fig. 5E).

**Figure 5.**
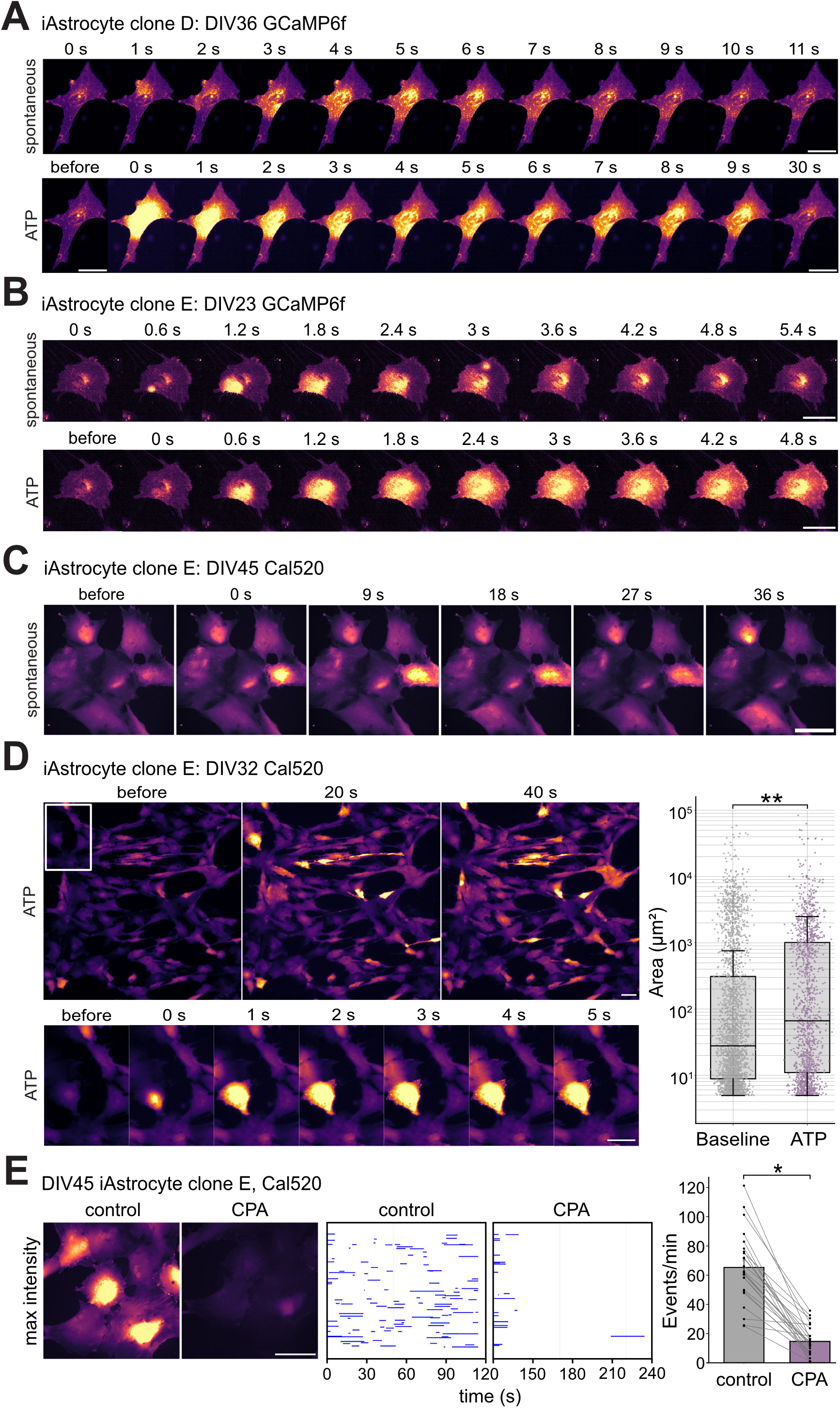
iAstrocytes have spontaneous calcium waves and react to ATP and CPA. **A)** Time course of a spontaneous calcium wave in DIV36 clone D derived iAstrocytes transduced with a GFAP promoter-driven membrane targeted GCaMP6f (upper panels), and calcium events evoked by 100 µM ATP (lower panels). Scale bars = 50 µm. **B)** Time course of a spontaneous calcium wave in DIV23 clone E derived iAstrocytes transduced with a GFAP promoter-driven membrane targeted GCaMP6f (upper panels), and calcium events evoked by 100 µM ATP (lower panels). Scale bars = 50 µm. **C)** Spontaneous calcium wave in DIV45 clone E derived iAstrocytes loaded with bath-applied calcium dye Cal520. Scale bar = 100 µm. **D)** Calcium response in DIV32 clone E derived iAstrocytes loaded with Cal520 and treated with 100 µM ATP (zoom of area indicated by white square is shown in lower panels). Scale bars = 50 µm. Quantitation of the area of calcium events detected by AQuA software in control conditions and following ATP application is shown on the right. **E)** DIV45 iAstrocyte clone E Cal520 calcium response to CPA. Scale bar = 100 µm. Asterisks indicate statistical significance (** p <0.001. * p <0.05; Student’s t-test).

### iAstrocytes support function and network activity of iNeurons

To test if iAstrocytes support iNeuron network formation, function and activity, we compared iNeurons alone, iNeurons with iAstrocytes (clone E), and iNeurons with mouse astrocytes in high-density multi-electrode array (HD-MEA) recordings. HD-MEAs were double-coated with PDL/Biolaminin, and iNeurons were plated. One week later, a five-minute-recording was acquired, after which iAstrocytes or mouse astrocytes were added. Five-minute recordings were then made weekly to assess network activity.

We found that iNeurons supported by iAstrocytes or mouse astrocytes had an earlier onset of activity, by DIV14, than iNeurons alone, which showed activity by DIV21 (Fig. 6A). This evolved into synchronous activity by DIV21 in mouse astrocyte supported cultures and DIV28 in iAstrocyte supported cultures. iNeurons supported by mouse astrocytes had higher activity that developed faster than iNeurons supported by iAstrocytes. This higher activity may mean that mouse astrocytes are superior to iAstrocytes in supporting iNeuron activity. However, it could also mean that mouse astrocytes cause aberrantly high activity in human neurons that represents a non-physiological state. iNeurons co-cultured with mouse astrocytes had earlier onset of network bursts, and a higher network burst frequency than iNeurons co-cultured with iAstrocytes, or iNeurons alone, which showed the most delayed increase in network burst frequency (Fig. 6B). Network burst duration was higher in iNeurons co-cultured with iAstrocytes, and in iNeurons alone, than in iNeurons co-cultured with mouse astrocytes (Fig. 6C). Mouse astrocytes may therefore increase network burst frequency and reduce burst duration of human neurons. The inter-burst interval was longer in iNeurons co-cultured with iAstrocytes and in iNeurons alone, than in iNeurons supported by mouse astrocytes (Fig. 6D), but the number of spikes per network burst was not different between the three conditions. We also observed lower coverage of the HD-MEA by iNeurons supported by iAstrocytes compared to mouse astrocytes. This may have contributed to the lower mean firing rate in iNeurons supported by iAstrocytes due to a lower number of connected neurons. It is also possible that the earlier participation of more electrodes with higher activity in the presence of mouse astrocytes may be because mouse astrocytes were more mature than iAstrocytes, at the time they were added to iNeurons.

**Figure 6.**
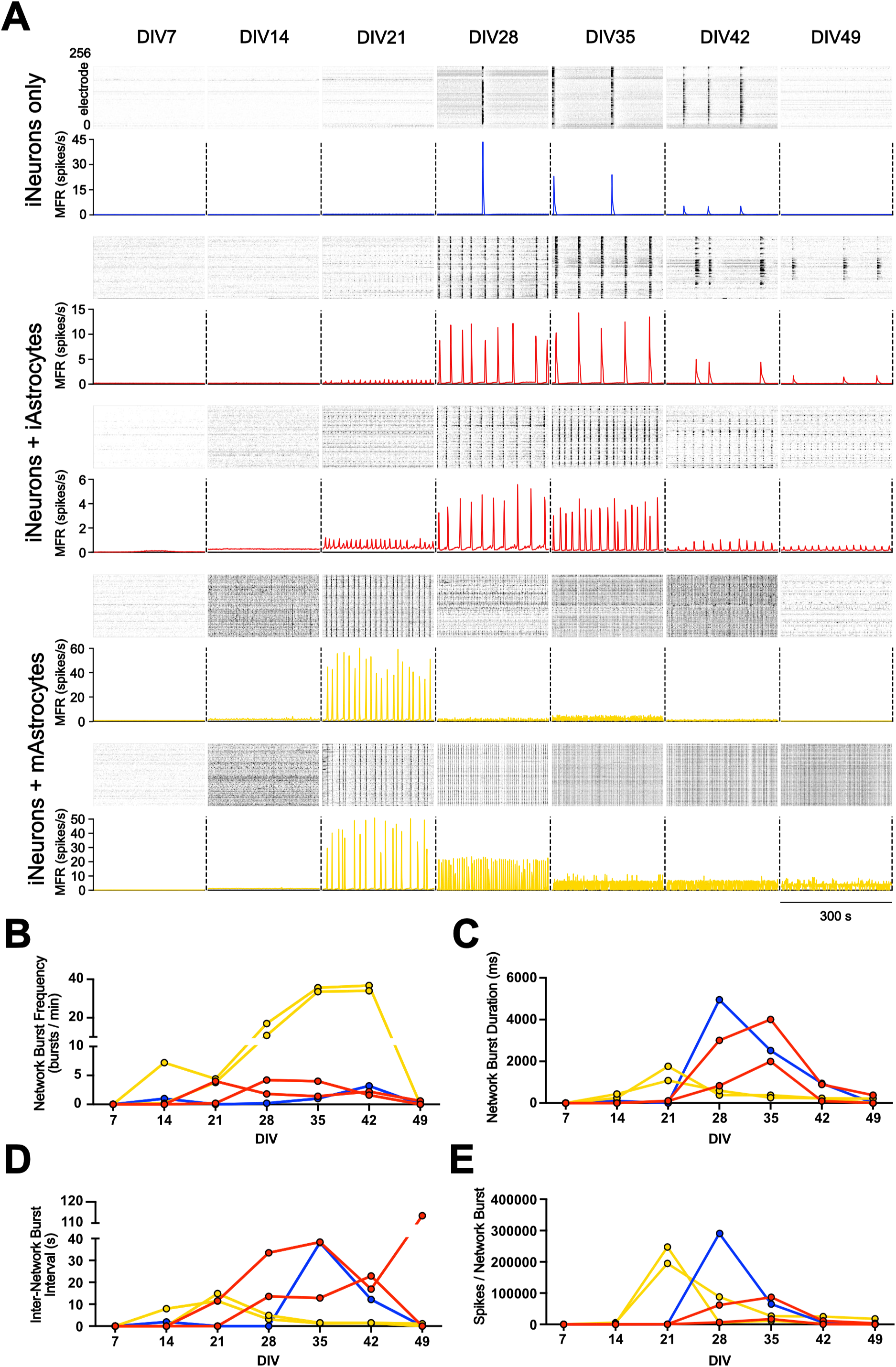
iAstrocytes support network activity of iNeurons grown on high-density multi-electrode arrays. **A)** Quantitation of network activity over time, recorded weekly, in iNeurons alone, iNeurons with iAstrocytes, and iNeurons with mouse astrocytes on 4096-channel HD-MEAs. Top panels are raster plots of activity from a randomly selected subset of 256 active electrodes, from weekly recordings five minutes in duration. The mean firing rate plots (bottom panels) show the total number of spikes detected per binning interval (1 second) divided by the total number (union) of all active electrodes on DIV7, 14, 21, 28, 35, and 49 combined. Technical replicates are shown separately, to highlight similarities and variability. **B)** Quantitation of network burst frequency, network burst duration **(C)**, inter-network burst interval **(D)**, and spikes/network burst **(E)** in weekly five-minute recordings over seven weeks, from DIV7 to DIV49.

### Calcium imaging confirms that iAstrocytes support iNeuron activity

We also tested network function in older (DIV56-61) cultures by calcium imaging of spontaneous activity in iNeurons supported by iAstrocytes, compared to iNeurons supported by mouse astrocytes. Calbryte 590 AM calcium dye was bath-applied to cultures, followed by imaging and analysis by Suite2P software, which can be used to extract fluorescence traces of individual cells that correspond to neurons. Raw fluorescence was then deconvolved into predictions of underlying action potentials by Cascade. DIV61 iNeurons supported by iAstrocytes showed robust mature synchronous activity (Fig. 7A), but with fewer network bursts than iNeurons co-cultured with mouse astrocytes (Fig. 7B), like the results of HD-MEA recordings. Putative iAstrocytes and mouse astrocytes (cells determined to be non-neuronal by Suite2P analysis) in co-cultures also showed spontaneous calcium activity.

**Figure 7.**
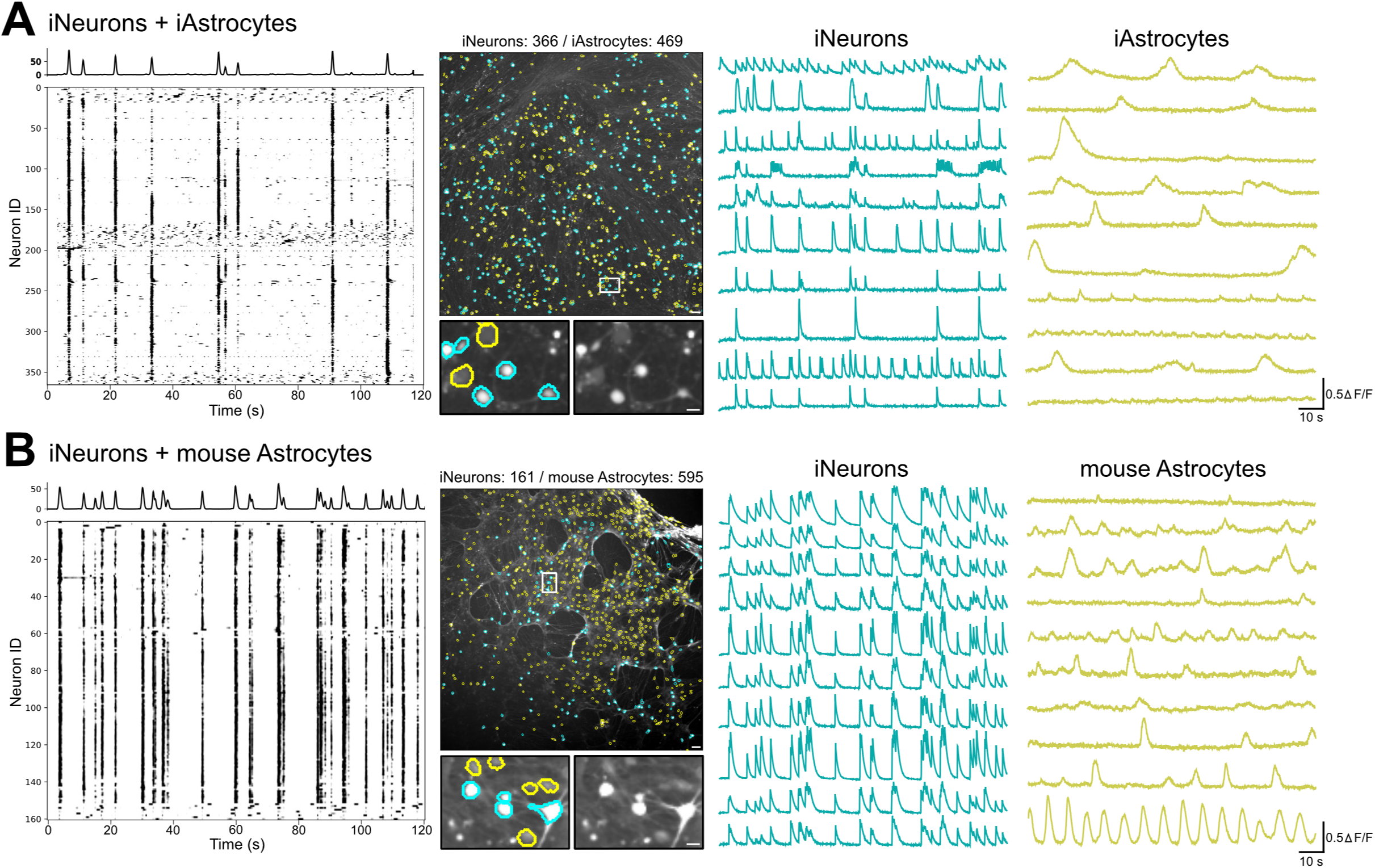
iAstrocytes support network activity assayed by calcium imaging. **A)** Raster plots of spontaneous calcium activity (left panel) in DIV61 iNeuron/iAstrocyte co-cultures loaded with Cal520, in neurons detected by Suite2P software (middle-left panel). Spike prediction from Cascade software is shown above the raster plot. Middle-left panel shows iNeurons (cyan) and putative iAstrocytes (yellow) detected by Suite2P software. Sample traces from individual neurons (cyan) and iAstrocytes (yellow) are shown on the right. **B)** Raster plots and spike prediction of spontaneous calcium activity in iNeurons co-cultured with mouse astrocytes. Middle-left panel shows iNeurons (cyan) and putative mouse astrocytes (yellow) detected by Suite2P software, and right panels show sample traces from individual neurons (cyan) and mouse astrocytes (yellow). Scale bars = 100 µm in top panels, and 25 µm in zooms below.

### iAstrocytes support iNeuron networks and synapses for at least 10 weeks

To test how long iAstrocyte-supported iNeuron networks were viable, we examined synapses and astrocyte and neuron morphology in immunostains of co-cultured networks grown for 71 days. DIV71 iAstrocyte (clone E)-supported iNeurons maintained numerous synapses, identified by synaptophysin immunostaining along MAP2 positive processes (Fig. 8A), like mouse astrocyte (mAstrocyte) and iNeuron co-cultures (Fig. 8B). iAstrocyte/iNeuron co-cultures also showed pronounced S100B and GFAP-positive astrocytes with stellate morphology and long processes in the vicinity of iNeurons (Fig. 8C). This suggests that iAstrocytes can be used to support iNeurons from at least three to ten weeks in culture.

**Figure 8.**
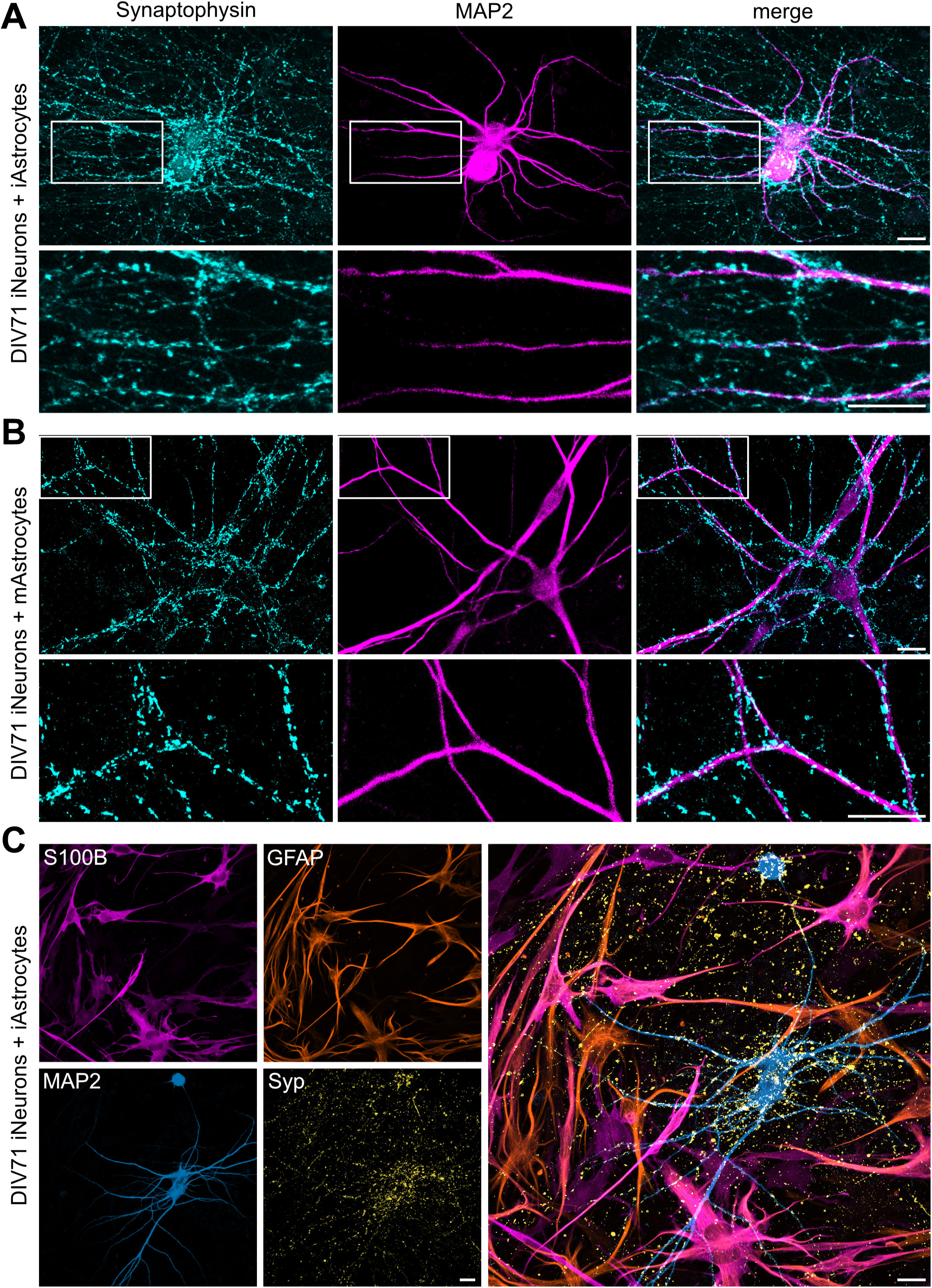
iAstrocytes support iNeuron networks for up to 71 days in culture. **A)** DIV71 iNeurons co-cultured with iAstrocytes immunostained with synaptophysin to mark synapses (cyan) and MAP2 to mark dendrites (magenta). **B)** DIV71 iNeurons co-cultured with mouse astrocytes, immunostained with synaptophysin (cyan) and MAP2 (magenta). **C)** DIV71 iNeuron/iAstrocyte co-cultures immunostained for S100B (magenta), GFAP (orange), MAP2 (blue) and synaptophysin (yellow). Scale bars = 20 µm.

## Discussion

We created and validated a stable, inducible SOX9/NFIB hiPSC line that produced functional mature human astrocytes (iAstrocytes). We further found that iAstrocytes support iNeuron growth, synapse formation, and network function and maturation.

Differential gene expression analysis of iAstrocytes, confirmed the upregulation and stable expression of NFIB and SOX9 expression over time. Gene expression during induction evolved from markers of the undifferentiated state of hPSCs to astrocytic markers, with the astrocytic markers S100B, PAX6, FABP7, GFAP, ALDH1L1, and SLC1A2 upregulated over time. Among these markers, S100B and SLC1A2 are expressed in mature astrocytes, indicating that iAstrocytes reach a mature state within a few weeks. Interestingly, AQP3, which is present in some astrocytes but is not an astrocyte-specific marker, was upregulated by induction, while the astrocytic-specific marker AQP4 was down-regulated. This may be due to the concomitant down-regulation of NFIA we observe in iAstrocytes, since NFIA/NFIB/SOX9 (and not NFIB/SOX9) upregulated AQP4 in transcription factor-induced human astrocytes (Baranes et al., 2023). The down-regulation of NFIA in our iAstrocytes is also surprising, given that SOX9 upregulates NFIA and NFIB (Deneen et al., 2006; Kang et al., 2012). Although NFIA expression is not shown in previous studies that used SOX9 to induce astrocytes, SOX9/NFIA expression only slightly increases NFIB. NFIA and NFIB might therefore be somewhat redundant in development of astrocytic fate (Bunt et al., 2017) where overexpression of one or the other is sufficient to induce mature astrocytes. Expression of SOX9 and NFIB in combination may therefore result in no change, or a downregulation of NFIA, as seen in our iAstrocytes. The glutamate transporter SLC1A2 was also unexpectedly downregulated, but this may be because expression of SLC1A2, as well as AQP4, is increased in co-cultures with neurons (Hasel et al., 2017). The levels of the common astrocytic markers GFAP and S100B were also markedly different in mouse versus human induced iAstrocytes. Mouse astrocytes had more GFAP, and iAstrocytes had more S100B and low levels of GFAP at three weeks in culture, possibly indicating differences in astrocyte reactivity in human iAstrocytes at this stage.

Our main goal was to test the ability of iAstrocytes to support human neuron growth and function. iAstrocytes specifically supported the growth of neurons; iNeurons preferentially grew on iAstrocytes and not on neighboring islands of undifferentiated progenitor cells. iAstrocytes also supported the formation of synapses with mature pre- and post-synaptic sites marked by synaptic vesicle markers and post-synaptic scaffolding proteins, by 28 days, with a similar number of synapses compared to cultures supported by mouse astrocytes, and significantly more than iNeuron cultures alone. This structural maturation was mirrored by function of human neuron networks, where neuronal firing became evident in iAstrocyte and mouse astrocyte supported iNeuron co-cultures by day 21-28 in HD-MEA recordings. We observed earlier and more synchronized firing in co-cultures of iNeurons and astrocytes than in iNeurons alone. iNeuron networks supported by mouse astrocytes had much higher activity that began earlier than iAstro-supported networks and showed more frequent synchronous activity. However, a higher mean firing rate and more network bursts may not necessarily indicate a “better”, more physiological network. It could rather represent a hyperexcitable network and pathological state. For example, human ALS-patient derived motor neurons have an increased mean firing rate in MEA recordings (Wainger et al., 2014). A preclinical mouse model of SYNGAP1-related intellectual disability causes hyperexcitability that increases the frequency of network bursts (Fenton et al., 2024). hiPSC-derived neuronal cultures with Alzheimer’s disease-related mutations in PSEN1 or APP have significantly higher mean firing rate and number of network bursts compared to isogenic gene corrected controls (Ghatak et al., 2019). Patient-derived networks with SCN1A mutations, which cause Dravet syndrome, a severe epileptic encephalopathy have increased mean firing rate and network burst frequency in response to proconvulsive compounds in MEA recordings (van Hugte et al., 2023). Thus, a higher mean firing rate and network burst frequency, as seen in mouse astrocyte supported human neuronal networks, may not necessarily represent a “better” electrophysiological phenotype. Although many early studies found more activity in human networks supported by rodent cultures than by human astrocytes (Lischka et al., 2018; Rhee et al., 2019) a few recent studies have found higher activity in cultures supported by human astrocytes than by rodent astrocytes (Lendemeijer et al., 2024; Neyrinck et al., 2021). These differences may be a result of different protocols to produce human astrocytes, as well as their maturation state, since human astrocytes are known to take longer to mature than rodent astrocytes. Human glial progenitor cells transplanted into the brains of neonatal mice have been shown to enhance long-term potentiation (LTP) and learning one year after transplantation (Han et al., 2013) suggesting that they may have superior capabilities given enough time to mature. The successful maintenance of iNeuron/iAstrocyte co-cultures to DIV 71 demonstrates that these cells can survive long-term.

Calcium imaging in older neurons (DIV61) showed a similar higher rate of synchronous firing in mouse astrocyte supported networks than in human iAstrocyte supported networks, but both iAstrocytes and mouse astrocytes supported mature synchronous activity. Support of networks by astrocytes (either mouse or human) does therefore seem to sustain normal network function.

We found that combining iNeuron Medium with iAstrocyte Maturation Medium and exchanging DMEM/F12 with advanced DMEM/F12 favored survival and maturation of iNeuron/ iAstrocyte co-cultures. In the future, iMicroglia could be added to this system, with the addition of the two factors IL-34 and GM-CSF to support the simultaneous maturation of microglia (which we have tested in preliminary experiments, not shown). The iNeuron Medium was modified from the original protocol (Canals et al., 2018) by adding ascorbic acid and using Neurobasal Plus and B27 Plus, rather than normal Neurobasal and B27. We also added the cell-permeable γ-secretase inhibitor Compound E, a Notch pathway inhibitor that promotes neuronal differentiation (Li et al., 2011). Other studies have used DAPT, another γ-secretase inhibitor for similar purposes (Meijer et al., 2019; Rhee et al., 2019).

Several protocols now exist to generate human astrocytes from hiPSCs. Stepwise differentiation of astrocytes by small molecules can be inefficient and time-consuming, but 90-day protocols (Krencik and Zhang, 2011) have now been reduced to as little as 28 days (Lendemeijer et al., 2024). Small molecule induction avoids introducing viruses or genetic modifications, and maintains more of the cell origin’s epigenetic marks, but results can be quite variable. Human astrocytes induced by transcription factors have the advantage that the technique is fast, efficient, reproducible and comparable across studies, and well-characterized in terms of mechanism and affected pathways. A potential loss of epigenetic age during reprogramming can obscure disease-relevant phenotypes but may also reveal early pathogenic changes before clinical onset. Our system could be particularly advantageous to standardize robust disease models. Direct conversion of fibroblasts into astrocytes with lentiviral SOX9/NFIB to preserve age and genetic heterogeneity of patients has been achieved, for example (Quist et al., 2022). Our gene-edited inducible hiPSC line provides human astrocytes that support mature human synapse and neuronal network formation in four weeks without the need to use lentivirus. This is an important advance to standardize the use of human astrocytes in neural networks and to advance translation of basic research to new clinical treatments of brain diseases and disorders. In the future, this culture system could be expanded to incorporate hiPSC-derived microglia, oligodendrocytes and inhibitory neurons in a complex network more closely representing neural circuits in the brain where disease mechanisms in diverse cell types could be investigated.

## Methods

### Generation of an inducible human iPSC line for production of iAstrocytes

#### NFIB/SOX9 plasmid construction

Two gene-edited hiPSC lines were initially made in which NFIA/SOX9 or NFIB/SOX9 was integrated into the AAVS1 locus of the BIHi005-A hiPSC line. The plasmid for NFIA/SOX9 (Li et al., 2018) was acquired from Addgene (AAVS1-TRE3G-NFIA+SOX9, Addgene 129455 from Su-Chun Zhang). To create the NFIB/SOX9 donor plasmid for gene editing, NFIB was amplified by PCR using Q5 High fidelity polymerase (M0492 S, NEB without a stop codon, from the tetO.Nfib.Hygro plasmid (Addgene 117271 from Henrik Ahlenius) introducing SalI and ClaI restriction site sticky ends in the primers: <u>SalI</u>-<u>Nfib</u>-fwd: CGTAAA<u>GTCGAC</u>GCCACC<u>atgatgtattctcccatctgtctcactcagg</u>, <u>ClaI</u>-NoStop-<u>Nfib1</u>-rev: CTTCC<u>ATCGATgcccaggtaccaggactggc.</u> The PCR product was double digested with ClaI and SalI (R0197/R3138 NEB) in rCutSmart Buffer (B6004S NEB) and a 1.2 kb band of the expected size was gel-purified. The vector plasmid AAVS1-TRE3G-NFIA+SOX9 was double digested with ClaI + SalI in rCutSmart Buffer and dephosphorylated with addition of a phosphatase for 20 minutes. The 12.7 kb (backbone) product was purified by agarose gel electrophoresis and the NFIB insert and AAVS1-TRE3G-SOX9 vector backbone ligated to form the new plasmid AAVS1-TRE3G-NFIB-SOX9.

#### Transfection of hiPSCs with plasmids for gene editing

Plasmids pZFN-AAV1-L-ELD (Addgene 159297) and PZFN-AAV1-R-KKR (Addgene 159298) from Kosuke Yusa (plasmid origin: Cambridge University, Dr. Mark Kotter) were used as left and right ZFN templates targeting the AAVS1 locus. BIHi005-A hiPSCs (https://hpscreg.eu/cell-line/BIHi005-A) were seeded as single cells at a density of 0.25x10^6^ cells/well of a 6-well plate in E8 with 10 µM Y-27632 ROCK inhibitor. hiPSC handling and culture was performed as previously described (Vallone et al., 2025). The day before transfection, hiPSCs at passage 40 were transfected with 3 µg NFIA-SOX9 or NFIB-SOX9 and 1 µg each of pZFN-AAV1-L-ELD and PZFN-AAV1-R-KKR plasmids. Two tubes containing 125 µl Opti-MEM (51985034 Thermo Fisher) were prepared. Plasmids (total 5 µg DNA) were added to one tube and 5 µl Lipofectamine Stem Transfection Reagent (STEM00001 Thermo Fisher) was added to the other. The solutions were incubated separately for 5 min, then mixed and incubated for another 10 min and then added to the hiPSCs after a medium change in E8 medium, and incubated for 30 hours at 32°C. Three days after transfection, the cells were selected with 250 ng/ml Puromycin (P8833 Sigma-Aldrich) for 6 days, collected, frozen with Bambanker serum-free cryopreservation media (Nippon Genetics BB01), and stored at -80°C.

#### Isolation and verification of gene-edited clones

One cryovial each of gene-edited hiPSCs (BIHi005-A_NFIA-SOX9 or BIHi005-A_NFIB-SOX9) were thawed and plated into two wells of a 6-well plate for each line. Cells were grown in 250 ng/ml Puromycin (P8833 Sigma-Aldrich) for selection in E8 medium (A1517001 Thermo Fisher) with Pen/Strep for three days, after which cells were split using 0.5 mM EDTA (Thermo Fisher AM9260G) and plated in StemFlex Medium (Thermo Fisher A3349401) with 1:20 CloneR2 (100-0691 StemCell Technologies). The next day, cells were dissociated into a single-cell suspension with StemPro Accutase (A11105-01 Thermo Fisher). Single hiPSC seeding and clone expansion was performed using isoCell (iotaSciences, IOTA-10010) as previously described (Vallone et al., 2020). Thirty-two clones per line were transferred to two 96 well plates coated with Geltrex (A1413302 Thermo Fisher), cultured until 60-80% confluent and frozen until DNA extraction and genotyping.

For genotyping DNA was extracted using the Phire Animal Tissue Direct PCR Kit (F140WH Thermo Fisher) following the manufacturer’s instructions. Sixteen clones per construct were screened with primer pairs (Table 1), testing for correct left and right homology arm insertion, NFIA and NFIB, and whether clones were homo- or heterozygous.

**Table 1.**
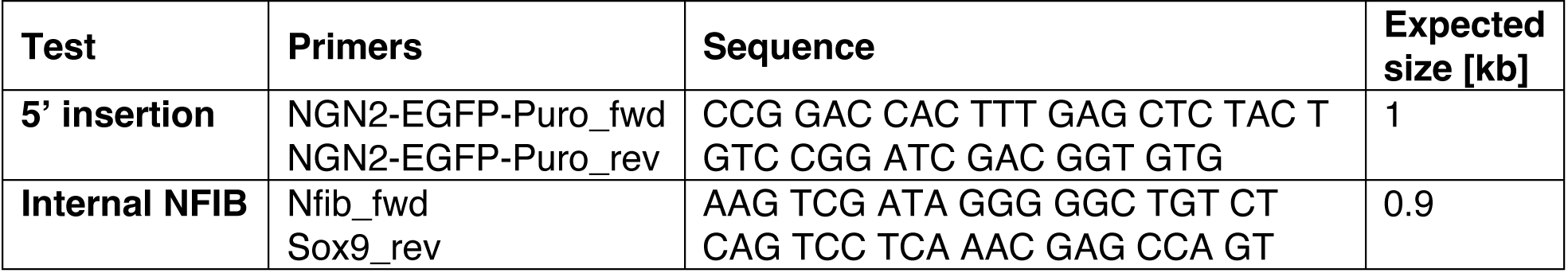
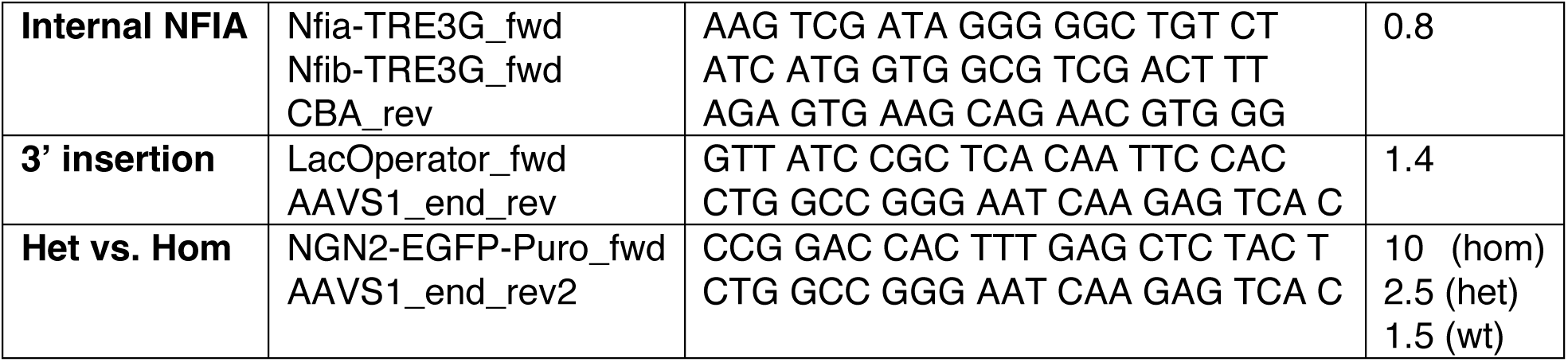
Primer pairs for genotyping NFIA-SOX9 and NFIB-SOX9 hiPSC clones.

For PCR, 25 ng DNA template, 10 µM each of forward and reverse primer, and 12.5 µl Q5 High fidelity Polymerase master mix (M0492S NEB) was added per 25 µl reaction. PCR program: 98 °C, 30 s; 35 x 98 °C 10s, 68/65 °C 15s, 72 °C 30s/1min/5min; 72 °C 2min; 6 °C inf. A GFP plasmid control, no template control, the parental line BIHi005-A, and another hiPSC reference line BIHi001-B, served as PCR controls. The NFIA-SOX9 line clones did not differentiate and mature to astrocytes reliably, so this line was not pursued further. We therefore focused on the NFIB/SOX9 hiPSC line, which has been reported to be superior to NFIB, NFIA, NFIA/SOX9, or NFIA/NFIB/SOX9 transcription factors in inducing astrocytic fate of hiPSCs (Canals et al., 2018). From NFIB-SOX9 editing, three clones, BIHi005-A-1C (homozygous), BIHi005-A-1D (heterozygous), and BIHi005-A-1E B5 (homozygous) (right graph) showed superior growth after editing and were banked for further analysis.

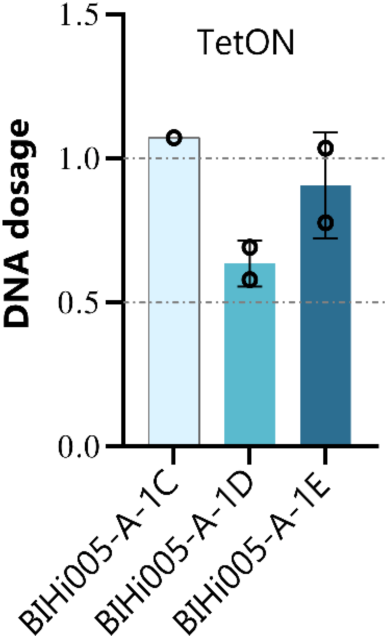

### Expansion and Differentiation of NFIB-SOX9 iAstrocytes

Cryovials of hiPSCs were quick-thawed by swirling vials in a water or bead bath and quickly transferred into 15 ml tubes containing 10 ml E8 medium (A1517001 Thermo Fisher) with 10 µM ROCK inhibitor (72305 StemCell Technologies), and centrifuged at 400 x g for 4 min at RT. The supernatant was removed, and cell pellet resuspended in E8 with 10 µM ROCK inhibitor and 3 µg/ml doxcycycline. Live cell number was determined using Trypan Blue Solution. Cells were plated at a density of 100,000 cells per well of a 6-well plate or 0.5 - 1.0 x 10^6^ cells per T75 cell culture flask coated with 5 µg/ml Vitronectin (A14700 Thermo Fisher Scientific) in sterile DPBS with Ca^2+^/Mg^2+^ for ≥ 1 h at 37°C. The first 8 days of differentiation follow the protocol published by Canals (2018). The day after plating (on Day of Induction, DoI, 1), the medium was fully exchanged with Expansion Medium (Table 2) containing doxycycline. On DoI 3 the transition to FGF Medium (Table 2) was begun by replacing Expansion Medium with a 3:1 mixture of Expansion Medium to FGF Medium. The next day, on DoI 4, medium was replaced with a 1:1 mixture of Expansion Medium: FGF Medium. On DoI 5, medium was replaced with a 1:3 mixture of Expansion Medium:FGF Medium. From DoI 6 onward, daily full-medium changes were continued with FGF Medium. At DoI 4, iAstrocytes usually needed to be split and were re-plated at a density of 100,000 cells per 6-well plate well or 1.0 x 10^6^ per T75 cell culture flask (10,500 – 13,500 cells/cm^2^). Every splitting was performed using StemPro® Accutase® (A11105-01, Lot 2442665 Thermo Fisher) and culture medium supplemented with 10 µM ROCK inhibitor (72305 StemCell Technologies). On DoI 7, 8 or 9 iAstrocytes were then frozen in Bambanker serum-free cryopreservation media (BB01, Nippon Genetics), added to cultured iNeurons, or plated alone for monocultures. For iAstrocyte monocultures half-medium changes were performed from DoI 10 onwards, as described in the original protocol (Canals et al. 2018), with Maturation Medium (Table 2).

**Table 2.** Media for Expansion and Differentiation of NFIB-SOX9 iAstrocytes.

| **Component** | **Final conc.** | **Cat. #** | **Source** |
| --- | --- | --- | --- |
| DMEM/F12 | 1x | 31330038 | Thermo Fisher |
| FBS | 10% | A5256801 | Thermo Fisher |
| N2 Supplement | 1% | 17502048 | Thermo Fisher |
| GlutaMAX | 1% | 35050038 | Thermo Fisher |
| Doxycycline* | 3 µg/ml | D3447 | Sigma-Aldrich |

***FGF Medium***
| **Component** | **Final conc.** | **Cat. #** | **Source** |
| --- | --- | --- | --- |
| Neurobasal | 1x | 21103049 | Thermo Fisher |
| B27 Supplement | 2% | 17504044 | Thermo Fisher |
| NEAA | 1% | 11140050 | Thermo Fisher |
| GlutaMAX | 1% | 35050038 | Thermo Fisher |
| FBS | 1% | A5256801 | Thermo Fisher |
| FGF (rec. human) | 8 ng/ml | 100-18B-10 | PeproTech |
| CNTF (rec. human) | 5 ng/ml | 450-13-20 | PeproTech |
| BMP4 (rec. human) | 10 ng/ml | 120-05ET-10 | PeproTech |
| Doxycycline* | 3 µg/ml | D3447 | Sigma-Aldrich |

***Maturation Medium***
| **Component** | **Final conc.** | **Cat. #** | **Source** |
| --- | --- | --- | --- |
| DMEM/F12 | 59% | 11330032 | Thermo Fisher |
| Neurobasal | 50% | 21103049 | Thermo Fisher |
| N2 | 1% | 17502048 | Thermo Fisher |
| Sodium Pyruvate | 1% | 35050038 | Thermo Fisher |
| GlutaMax | 1% | 35050038 | Thermo Fisher |
| N-acetyl-cysteine | 5 µg/ml | A8199 | Sigma-Aldrich |
| EGF-like growth factor, heparin-binding (human) | 5 ng/ml | 130-093-825 | Miltenyi Biotec |
| CNTF (rec. human) | 10 ng/ml | 450-13-20 | PeproTech |
| BMP4 (rec. human) | 10 ng/ml | 120-05ET-10 | PeproTech |
| dbcAMP | 500 µg/ml | D0627 | Sigma-Aldrich |

\*added fresh directly to each aliquot for medium change.

### RNA Sequencing of iAstrocytes

At day of induction (DoI) 0 (before Doxycycline induction), and at DoI 3, 6, 13, 21 and 27 iAstrocyte cell lysates were collected in triplicate and stored frozen in Lysis Solution. RNA was extracted using the Aurum Total RNA Mini Kit (Bio-Rad 7326820). Identical control samples, in which no doxycycline was applied, were collected in duplicate until day 13 (after which culture health declined). RNA quality and concentration were evaluated by Tape Station (Agilent) and NanoDrop, respectively. Next generation sequencing was performed by Azenta Life Sciences. We then analyzed gene expression with R Project for Statistical Computing (version 4.4.2, RRID:SCR_001905) by combining the R packages DESeq2, RRID:SCR_015687 (Love et al. 2014), clusterProfiler, v4.14.4, RRID:SCR_016884 (Xu et al. 2024), ComplexHeatmap version 2.22.0, RRID:SCR_017270 (Gu et al. 2016), enrichplot v1.26.6, a gene over-representation analysis with MeSH term enrichment analysis (Yu 2018) and CellMarker 2.0 (Hu et al. 2023). Ensembl (RRID:SCR_002344) was used for gene identification, UniProt (RRID:SCR_002380) for information on encoded proteins and their function.

### Preparation of mouse primary astrocytes

Mouse astrocytes were prepared from E18-E19 timed pregnant C57BL/6J mice, euthanized with CO_2_ and by cervical dislocation according to the specifications of the Institutional Animal Care and Ethics Committee of Charité University (T-CH 0013/20), and German animal welfare laws. The mouse was placed on its back, the abdomen disinfected with 70% ethanol, and skin and muscles dissected to expose the uterus. Embryos were extracted from amniotic sacs and decapitated. Heads were transferred to a 10 cm petri dish filled with ice-cold dissection solution: HBSS (Gibco Hanks’ Balanced Salt Solution, without Ca^2+^/Mg^2+^, 14170138 Thermo Fisher) and 10 mM HEPES (15630080 Thermo Fisher). Heads were fixed ventral side down with forceps. After removing the cerebellum and back of the skull with small scissors, the hemispheres were extracted, meninges removed, hippocampus cut out, and cortices cut into 4-8 pieces and transferred into a 15 ml Falcon tube filled with ice-cold dissection solution. Dissection solution was replaced with 2-3 ml of 0.25% pre-warmed (37°C) Trypsin (25200056, Thermo Fisher), and incubated in a water bath for 20 min at 37°C. The tissue was then washed three times with 5 ml dissection solution, and cells triturated with a 1000 ul pipette tip in 2 ml mouse astrocyte culture medium: DMEM (high glucose, GlutaMAX Supplement, pyruvate 10569010, Thermo Fisher) with 1% Penicillin-Streptomycin (10.000 U/ml, 15140122 Thermo Fisher) and 10% FBS (F0804, Sigma-Aldrich/Merck). The cell suspension volume was increased to 10 ml and filtered through a 100 µm cell strainer. 1-2 million cells in 12 ml mouse astrocyte medium were plated per T75 flask coated with a collagen-polyornithine mix containing 0.12 µg/mL poly-DL-ornithine (P8638, Sigma-Aldrich), 0.66% (v/v) Collagen type I (C3867, Sigma), 0.06% (v/v) and acetic acid (Roth) in water (Schiweck et al., 2021) for 1-2 h at 37°C. After a full medium change the day after plating, medium changes were performed once a week until cells were split or used for co-culture. Astrocytes became confluent in 3-5 weeks. To split confluent astrocytes, medium was removed, astrocytes were washed with 10 ml pre-warmed HBSS (Gibco Hanks’ Balanced Salt Solution, without Ca^2+^/Mg^2+^, 14170138 Thermo Fisher) and incubated with 5 ml pre-warmed (37°C) 0.25% Trypsin-EDTA (25200056 Thermo Fisher) or 10x TrypLE Select (A12177-01 Thermo Fisher) for 5-10 min until cells detached from the surface. 5 ml medium was then added, and cells resuspended by gently tapping the culture vessel and rinsing the surface with medium. Cells were transferred to a 15 ml tube, centrifuged at 400 x g for 5 min at RT, resuspended in DMEM (10569010, Thermo Fisher), centrifuged again at 400 x g for 5 min at RT, and finally resuspended in the medium for co-culture with iNeurons, or in mouse astrocyte medium if astrocytes were split into a new T75 flask. Mouse astrocytes were grown for 4-8 weeks before adding them to iNeurons for co-cultures and were used at a maximum 8 weeks in culture, and 3 passages.

### iNeuron culture

The hiPSC line BIHi005-A-24 (https://hpscreg.eu/cell-line/BIHi005-A-24) was pre-differentiated in Neural Induction Medium (DMEM/F12 with 2.5 mM Glutamine, 15 mM HEPES, 1x B27, 1x N2-Supplement, 2 µM Dorsomorphin, 10 µM SB431542, 1x Pen/Strep) for one week to create neural stem cells (NSCs), which were frozen as a single batch in Bambanker serum-free cryopreservation media (BB05, Nippon Genetics), to ensure reproducibility in successive experiments.

NSCs were quick thawed and transferred into 15 ml tubes containing 10 ml Advanced DMEM/F12 supplemented with 10 µM Y-27632 ROCK inhibitor. Cells were centrifuged at 400 x g for 4 min at RT, and the cell pellet resuspended in Modified Neural Expansion Medium (Table 3). Live cells were determined by Trypan Blue exclusion and plated at a density of 90,000-100,000 cells per well of 24-well plates, coated with 0.5 mg/ml Poly-D-Lysine (P7886, Sigma-Aldrich) in 0.1 M borate buffer (0.31 g boric acid, B9645, Sigma-Aldrich, in 50 mL ddH_2_O, pH 8.5, 0.22 µm sterile-filtered) for ≥ 2 h or O/N at RT/37°C, or 5 µg/ml Vitronectin (A14700, Thermo Fisher) in sterile DPBS without Ca^2+^/Mg^2+^ for ≥ 1 h or O/N at 37°C, and then 2-10 µg/ml Bio-laminin-521 (LN521-05, Bio-lamina) in sterile DPBS with Ca^2+^/Mg^2+^ for 2-3 h at 37°C. Cells were then cultured in Neural Expansion Medium for 24 - 48 h at 37°C, 5% CO_2_. The day after plating, medium was changed to iNeuron medium (Table 3).

**Table 3.** Media for iNeuron culture.

***Modified Neural Expansion Medium***
| **Component** | **Final conc.** | **Cat. #** | **Source** |
| --- | --- | --- | --- |
| Neurobasal Medium | 0.5x | 21103049 | Thermo Fisher |
| Advanced DMEM/F-12 | 0.5x | 12634010 | Gibco/Thermo Fisher |
| B27 supplement | 1x | 17504044 | Thermo Fisher |
| Neural Induction Supplement | 1x | A16477-01 | Thermo Fisher |
| N2-Supplement | 1x | 17502048 | Gibco/Thermo Fisher |
| GlutaMAX Supplement | 1x | 35050038 | Gibco/Thermo Fisher |
| human rec. BDNF | 10 ng/ml | 450-02, Lot 122261, A04523 | PeproTech |
| human rec. NT-3 | 10 ng/ml | 267-N3-025 | R&D Systems |
| Laminin | 0.2 μg/ml | L2020 | Sigma-Aldrich |
| ROCK Inhibitor | 10 μM | 72305 | Stem Cell Technologies |
| Pen/Strep | 1x | 15140122 | Thermo Fisher |
| Doxycycline | 3 μg/ml | D3447 | Sigma-Aldrich |

***iNeuron Medium***
| **Component** | **Final conc.** | **Cat. #** | **Source** |
| --- | --- | --- | --- |
| Neurobasal/Plus Medium | 0.5x | 21103049/<br>A3582901 | Thermo Fisher |
| B27/Plus Supplement | 1x | 17504044/<br>A3582801 | Thermo Fisher |
| GlutaMAX Supplement | 1x | 35050038 | Gibco/Thermo Fisher |
| human rec. BDNF | 10 ng/ml | 450-02, Lot 122261, A04523 | PeproTech |
| human rec. NT-3 | 10 ng/ml | 267-N3-025 | R&D Systems |
| Laminin | 0.2 μg/ml | L2020 | Sigma-Aldrich |
| Compound E | 0.2 μM | 6476, Cas No 209986-17-4, Batch 4B/263805 | R&D Systems/<br>Tocris |
| Ascorbic acid 2-phosphate | 200 μM | A8960, Lot SLCN6315 | Sigma-Aldrich |
| CultureOne Supplement* | 1x | A33202-01, Lot 2463536 | Thermo Fisher |
| AraC** | 2 μM | C1768 | Sigma-Aldrich |
| Doxycycline | 3 μg/ml | D3447 | Sigma-Aldrich |

\* Can be added to inhibit neural progenitor overgrowth, if this is detected in the culture.
\*\* Added for 24 h. 2-4 days after adding astrocytes.

Daily full medium changes were then performed for the first 7 days of induction (DoI), with fresh 3 µg/ml doxycycline directly added to the medium aliquot each day. On DoI 2, mouse astrocytes were added for iNeuron/mouse Astrocyte co-cultures. Two days later, on DoI 4, 2 µM AraC (C1768 Sigma-Aldrich) was added for 24 h to inhibit astrocyte and progenitor overgrowth. After DoI 7, half-medium changes were performed continually every 2-3 days.

### Co-culture of iAstrocytes with iNeurons

Split or quick-thawed iAstrocytes were plated at a density of 50,000 cells per well of a 24-well plate well on iNeurons cultured until DoI 2-7 in Co-Culture Medium (Table 4), a 1:1 mix of iNeuron Medium (without ascorbic acid, CultureOne Supplement or AraC) and iAstrocyte Maturation Medium, supplemented with 3 µg/ml doxycycline, plus 10 µM ROCK inhibitor. After a full medium change to Co-Culture Medium the day after plating, half of the medium was changed every 2-3 days, using Co-Culture Medium always containing fresh doxycycline.

**Table 4.** Media for co-culture of iAstrocytes with iNeurons.

*Co-Culture Medium*
| Component | Final conc. | Cat. # | Source |
| --- | --- | --- | --- |
| Doxycycline | 3 μg/ml | D3447 | Sigma-Aldrich |
| Neurobasal Medium/Plus | 50% | 21103049/A3582901 | Thermo Fisher |
| Advanced DMEM/F-12 | 50% | 12634010 | Thermo Fisher |
| GlutaMAX | 1% | 35050038 | Thermo Fisher |
| B27 supplement/Plus | 1% | 17504044/A3582801 | Thermo Fisher |
| hBDNF (rec. human) | 10 ng/ml | 450-02, Lot 122261, A04523 | PeproTech |
| hNT3 (rec. human) | 10 ng/ml | 267-N3-025 | R&D Systems |
| Laminin | 0.2 μg/ml | L2020 | Sigma-Aldrich |
| Compound E (after 24 h) | 0.2 μM | 6476 (4B/263805) | Tocris |
| Ascorbic acid 2-phosphate | 200 μM | A8960, Lot SLCN6315 | Sigma-Aldrich |
| CultureOne Supplement* | 1% | A33202-01, Lot 2463536 | Thermo Fisher |
| N2 Supplement | 1% | 17502048 | Thermo Fisher |
| N-acetyl-cysteine | 5 μg/ml | A8199 | Sigma-Aldrich |
| hbEGF-like growth factor | 5 ng/ml | 130-093-825 | Miltenyi Biotec |
| CNTF (rec. human) | 10 ng/ml | 450-13-20 | PeproTech |
| BMP4 (rec. human) | 10 ng/ml | 120-05ET-10 | PeproTech |
| dbcAMP | 500 μg/ml | D0627 | Sigma-Aldrich |
| AraC (Cytosine arabinoside)** | 2 μM | C1768 | Sigma-Aldrich |

\* Added beginning on day 4 of co-culture to inhibit overgrowth of progenitor cells.
\*\*Added for 24 h on day 4 of co-culture to reduce astrocyte and progenitor growth.

### Immunofluorescence, imaging and analysis of fixed samples

Cells were fixed with 4% PFA (J61899-AK Thermo Fisher) for 15 min and washed three times with 1x PBS (10010023 Thermo Fisher). Antibody labelling was performed either by 1) Blocking unspecific binding in 2% donkey Serum (D9663 Sigma-Aldrich), 0.1% w/v Triton-X-100 (3051 Roth) in 2x PBS (phosphate-buffered saline pH 7.4; 70011044 Thermo Fisher) for 30-60 min at RT, followed by replacing blocking solution with the same solution containing primary antibodies, or 2) Blocking in 2.5% donkey serum, 2.5% goat serum (G6767 Sigma-Aldrich), and 1% Triton X-100 in 1x PBS for 30-60 min at RT, followed by replacing blocking solution with primary antibodies (Table 6) in 0.5% donkey serum, 0.5% goat serum, and 0.2% Triton X-100 in 1xPBS. Cells were incubated at 4°C O/N and then washed three times for 10 min with 1x PBS. Secondary antibodies (Table 6) were then added in the respective buffers and cells incubated for 2 h at RT on a shaker set to 18 rpm. Cells were washed three times 5 min each with 1x PBS. For DAPI stain, 1 μg/ mL DAPI in 1x PBS was added to cells for 15 min at RT on a shaker, and cells were washed three times 5 min each with 1x PBS. Coverslips mounted with Fluoromount-G (00-4958-02 Thermo Fisher) were allowed to dry and sealed with nail polish. Cells plated on 24-well glass bottom plates (with high performance #1.5 cover glass, P24-1.5H-N Cellvis) were kept in 1x PBS until imaged.

**Table 6.**
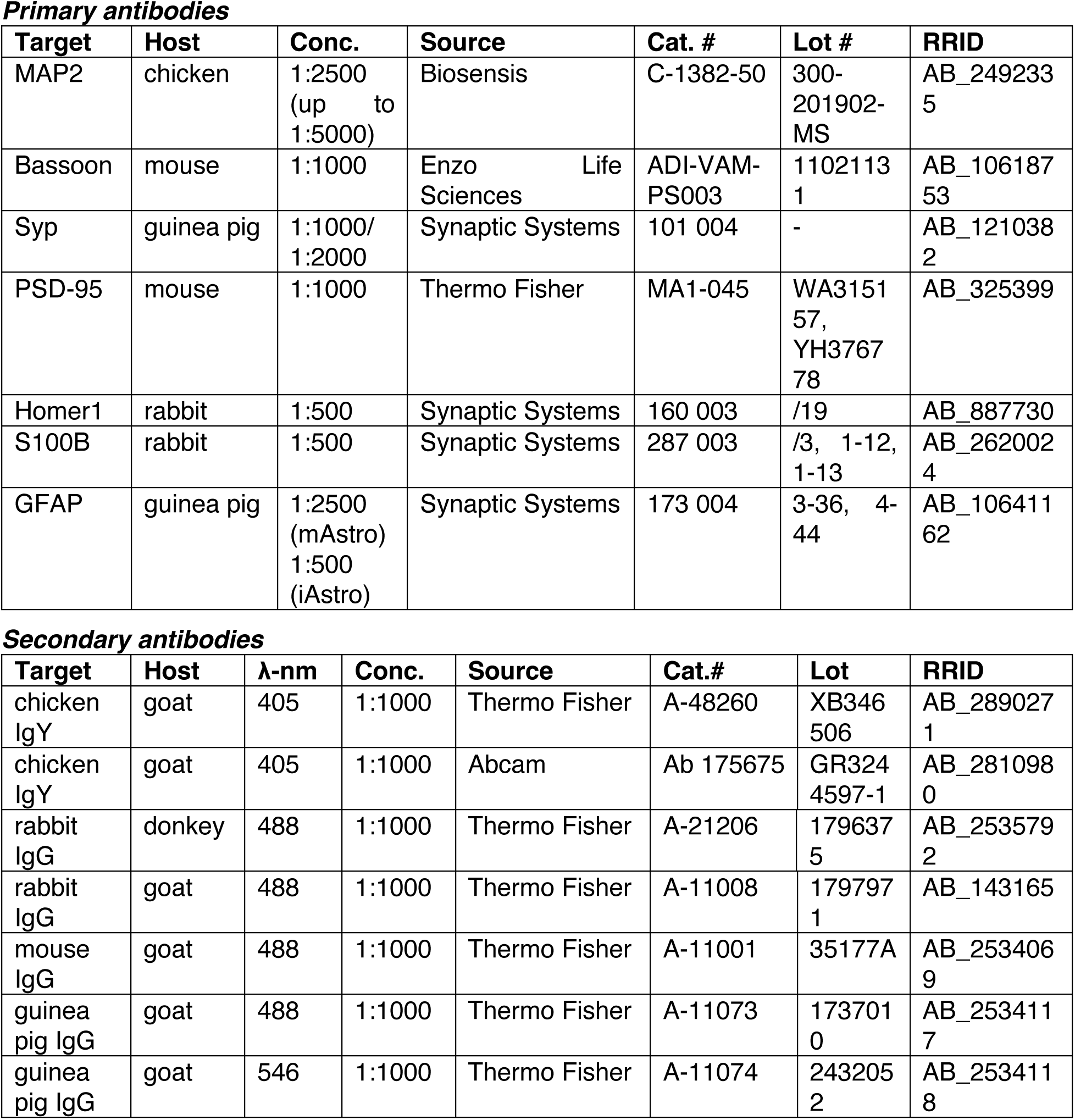
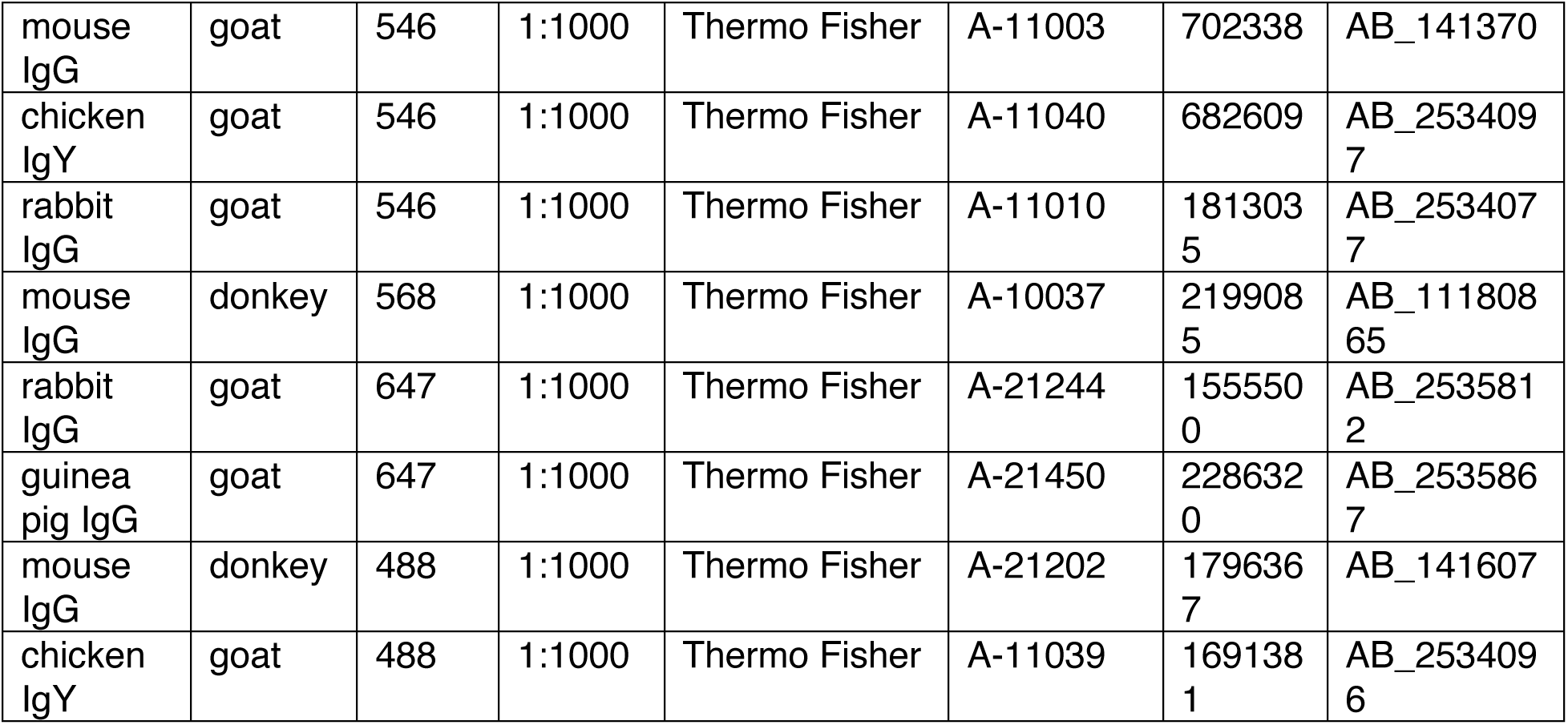
Primary and secondary antibodies.

| Target | Host | Conc. | Source | Cat. # | Lot # | RRID |
| --- | --- | --- | --- | --- | --- | --- |
| MAP2 | chicken | 1:2500<br>(up to<br>1:5000) | Biosensis | C-1382-50 | 300-201902-MS | AB_249233<br>5 |
| Bassoon | mouse | 1:1000 | Enzo Life<br>Sciences | ADI-VAM-PS003 | 1102113<br>1 | AB_106187<br>53 |
| Syp | guinea pig | 1:1000/<br>1:2000 | Synaptic Systems | 101 004 | - | AB_121038<br>2 |
| PSD-95 | mouse | 1:1000 | Thermo Fisher | MA1-045 | WA3151<br>57,<br>YH3767<br>78 | AB_325399 |
| Homer1 | rabbit | 1:500 | Synaptic Systems | 160 003 | /19 | AB_887730 |
| S100B | rabbit | 1:500 | Synaptic Systems | 287 003 | /3, 1-12,<br>1-13 | AB_262002<br>4 |
| GFAP | guinea pig | 1:2500<br>(mAstro)<br>1:500<br>(iAstro) | Synaptic Systems | 173 004 | 3-36, 4-44 | AB_106411<br>62 |

***Secondary antibodies***
| Target | Host | λ-nm | Conc. | Source | Cat.# | Lot | RRID |
| --- | --- | --- | --- | --- | --- | --- | --- |
| chicken<br>IgY | goat | 405 | 1:1000 | Thermo Fisher | A-48260 | XB346<br>506 | AB_289027<br>1 |
| chicken<br>IgY | goat | 405 | 1:1000 | Abcam | Ab 175675 | GR324<br>4597-1 | AB_281098<br>0 |
| rabbit<br>IgG | donkey | 488 | 1:1000 | Thermo Fisher | A-21206 | 179637<br>5 | AB_253579<br>2 |
| rabbit<br>IgG | goat | 488 | 1:1000 | Thermo Fisher | A-11008 | 179797<br>1 | AB_143165 |
| mouse<br>IgG | goat | 488 | 1:1000 | Thermo Fisher | A-11001 | 35177A | AB_253406<br>9 |
| guinea<br>pig IgG | goat | 488 | 1:1000 | Thermo Fisher | A-11073 | 173701<br>0 | AB_253411<br>7 |
| guinea<br>pig IgG | goat | 546 | 1:1000 | Thermo Fisher | A-11074 | 243205<br>2 | AB_253411<br>8 |
| mouse IgG | goat | 546 | 1:1000 | Thermo Fisher | A-11003 | 702338 | AB_141370 |
| chicken IgY | goat | 546 | 1:1000 | Thermo Fisher | A-11040 | 682609 | AB_2534097 |
| rabbit IgG | goat | 546 | 1:1000 | Thermo Fisher | A-11010 | 1813035 | AB_2534077 |
| mouse IgG | donkey | 568 | 1:1000 | Thermo Fisher | A-10037 | 2199085 | AB_11180865 |
| rabbit IgG | goat | 647 | 1:1000 | Thermo Fisher | A-21244 | 1555500 | AB_2535812 |
| guinea pig IgG | goat | 647 | 1:1000 | Thermo Fisher | A-21450 | 2286320 | AB_2535867 |
| mouse IgG | donkey | 488 | 1:1000 | Thermo Fisher | A-21202 | 1796367 | AB_141607 |
| chicken IgY | goat | 488 | 1:1000 | Thermo Fisher | A-11039 | 1691381 | AB_2534096 |

Images of fixed samples were acquired on a PerkinElmer Opera Phenix high content screening system, an SP-5 Leica DM 6000 upright confocal microscope with 405, 458, 476, 488, 496, 514, 561 and 633 nm laser lines using a 63x oil immersion objective, an SP-8 Leica DMI 6000 inverted confocal microscope with 405, 488, 552, 635 nm lasers using 40x and 63x oil immersion objectives, or a Nikon Spinning Disk Confocal CSU-W1 SoRa with 405, 445, 488, 515, 561, 594, and 638 nm laser lines and a 2x Hamamatsu ORCA-Fusion digital sCMOS camera (2304x2304 pixels). High content data acquired by the Opera Phenix was analyzed using Signals Image Artist (Revvity) software, to quantify cell-type specific markers and cell numbers. Images were first filtered by smoothing with a Gaussian width of 2-3 pixels to generate contiguous fluorescence signal of objects. Nuclei were determined by DAPI-labelled fluorescent objects < 500 µm^2^, > 70 µm^2^, with a roundness score > 0.85, empirically determined. A perfectly round object has a score of 1. Roundness is calculated by the equation:

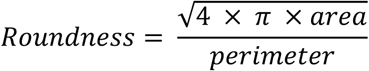

MAP2 signal above a fluorescence intensity threshold determined in control conditions without iNeurons was used to identify neurons with the Find Surrounding Region function using the Input Region of identified nuclei. MAP2 objects with average intensity < 5000 AU were excluded, since control wells without iNeurons had an average MAP2 intensity < 5000 AU. GFAP and S100B immunofluorescence were used to identify astrocytic cell populations using the Find Surrounding Region function with Input Region of nuclei.

Quantitation of synapses per length dendrite along MAP-2 positive processes was performed by counting Synaptophysin-positive puncta overlapping MAP2 signal in 20 µm lengths of dendrites (three random regions per image field) in a blinded fashion using FIJI/ImageJ Version 2.1.0/1.53c.

### Live imaging

#### iAstrocyte calcium imaging and analysis

iAstrocytes were transduced with AAV5 gfaABC1D-lck-GCaMP6f (GFAP promoter-driven membrane targeted GCaMP6f; Addgene 52924 from Baljit Khakh) with 0.4 µl of 7×10¹² vg/ml virus per well of a 24-well two weeks prior to imaging (subsequently diluted by half-medium changes every other day), or incubated with 2-5 µM Cal-520 AM (ab171868 Abcam) in 24-well or 96-well plates for 30-60 min. Cells in Gibco FluoroBrite DMEM (A1896701 Thermo Fisher) were imaged for 3 min baseline and 3 min more after addition of 50-100 µM ATP, at a 300 ms frame rate and 100 ms exposure time, with 2x2 pixel live binning to reduce file size and improve signal-to-noise, using an APO LWD 40x WI Delts objective and with 470 nm excitation and a GFP 579/26 nm emission filter on a Nikon Widefield Ti2 epifluorescence microscope with a sCMOS, PCO.edge camera and LED Illumination (Lumencore, SpectraX) and live-cell incubator at 37°C and 5% CO_2_. Timelapse recordings were analyzed with Astrocyte Quantitative Analysis (AQuA) software (Wang et al., 2019). 20-30 frames from the beginning of each recording were removed to improve signal detection. Area of calcium events was quantified as a continuous parameter in a linear mixed model with random intercept and slope. Spontaneous and ATP-evoked calcium event frequencies were compared using a non-parametric Wilcoxon signed rank test for paired data.

#### iNeuron calcium imaging and analysis

iNeurons plated on glass bottom 24-well plates (Cellvis) at a density of 80,000 cells per well, with 30,000 cells per well of clone D iAstrocytes or mouse Astrocytes added two days later, were incubated with 2 µM Calbryte 590 AM (20700, AAT Bioquest) for 20-30 min in media on the day of recording, and then imaged in modified ACSF (110 mM NaCl, 5 mM KCl, 1 mM NaH_2_PO_4_, 25 mM Glucose, 26 mM NaHCO_3_, 1 mM MgCl, 2 mM CaCl_2_, 10 mM HEPES, pH 7.2). Time-lapse images were acquired at 10 frames per second with 75 ms exposure for 2 minutes, with 2x2 pixel live binning to reduce file size and improve signal-to-noise, on a Nikon Eclipse Ti2 widefield epifluorescence microscope with a sCMOS, PCO.edge camera and LED Illumination (Lumencore, SpectraX) and live-cell incubator at 35°C and 5% CO_2_. Calcium imaging recordings were analyzed by a custom Python analysis pipeline that integrates cell detection with Suite2p (Pachitariu et al., 2017; v0.14.4, https://www.biorxiv.org/content/10.1101/061507v2) and deconvolution with Cascade v1.0.0 (Rupprecht et al., 2021). Suite2p and its sister cell detection algorithm, Cellpose (v3.1.0), are used to automatically detect fluorescent cells based on average fluorescence over the duration of a recording. Cascade then converts the raw fluorescence from each individual cell into predictions of underlying action potentials based on a database of paired calcium imaging and electrophysiological recordings. Potential ROIs are first filtered automatically based on their predicted activity. A minimum threshold of 0.1 predicted spikes, empirically determined by testing detection of active cells in the presence of 1 uM TTX, is set for an ROI to be considered ‘active’. The ‘active’ cells are next manually sorted based on their morphology and the shape of their fluorescence activity patterns into either neuronal or non-neuronal cells and noise, where neurons are bright and round with fast changes in fluorescence, and astrocytes are dim and oblong with slower changes in fluorescence. Individual traces are plotted using matplotlib (v3.10.0) while raster plots were created with the program Rastermap v1.0 (Stringer et al., 2025) using single-cell cascade predicted spikes and the total number of predicted spikes per frame across all cells. The code used for analysis is available upon request.

### Multielectrode array recordings and analysis

High-density Accura CorePlate1W 38/60 microelectrode arrays (3Brain) with a recording area of 3.8 mm x 3.8 mm containing 4096 electrodes, each 21 µm x 21 µm, in a 64 × 64 electrode-array, equally spaced with a 60 µm pitch were used on a BioCAM Duplex MEA system (3Brain). HD-MEAs were coated with 2 µg/ml poly-DL-ornithine hydro-bromide (P8638, Sigma-Aldrich) in ddH_2_O, sterile filtered, for 2 h at 37°, washed three times with sterile ddH_2_O, and coated with 10 µg/ml Bio-Laminin (LN521-05, Bio-lamina) diluted in sterile DPBS with Ca^2+^ and Mg^2+^ (14040-083 Gibco) for 2 h at 37°C. Bio-Laminin coating was carefully removed leaving a thin film on the chip such that it was not allowed to dry and iNeurons were plated immediately at a density of 75,000 cells in a 75 µl drop of NEM with 3 µg/ml Doxycycline and 10 µM ROCK inhibitor, on the MEA surface. After 1 h at 37°C when cells had attached to the MEA, the well volume was filled to 1.5 ml with NEM and maintained in iNeuron Medium with daily medium changes. Cultures were treated with 2 mM AraC for 24 h on day four to prevent proliferation of progenitor cells. One week later, BIHi-005-24-1D iAstrocytes were added at a density of 375,000 cells per well for iNeuron/iAstrocyte co-cultures. The same density of mouse astrocytes (DIV24, no split) was added for iNeuron/mouse Astrocyte co-cultures. Astrocytes were diluted in Co-culture Medium with 10 µM ROCK inhibitor and added to iNeurons during a full media change. The iNeuron only MEAs also received a full media change, but without iAstrocytes or mouse Astrocytes present in the Co-Culture Medium. The following day another full-media change was performed. Cultures were then maintained in Co-culture Medium with half media changes every 2-3 days. 2 µM AraC was added again for 24 h, four days after addition of astrocytes to prevent their overgrowth. MEAs were recorded every 7 days (at 37°C and 5% CO_2_) the day after a half media change to allow cultures to recover after feeding. BrainWave 5.6 software (3Brain) was used for acquisition, pre-and post-processing and analysis. Results were exported in csv format and plotted using Python v. 3.12.7. To construct raster plots .bxr files were further analyzed with Python (RRID:SCR_008394), packages h5py (RRID:SCR_024812), NumPy (RRID:SCR_008633),

Pandas (RRID:SCR_018214), MatPlotLib (RRID:SCR_008624), seaborn (RRID:SCR_018132).The mean firing rate was calculated by dividing the sum of all spikes detected in one second by the number of active electrodes. Active electrodes were defined as having ≥ 0.1 spikes/second during a five-minute recording session. Electrodes with ≤ 0.1 spikes/second were filtered out and considered noise, given that TTX application had no effect on spiking units below this threshold. Network bursts were defined as at least 10% of active units having a total of at least 50 spikes within 50 ms. Recordings were done at 37C° with humidified 5% CO_2_ perfusion via a mini chamber placed on top of the HD-MEA chip (Pecon® CO_2_-controller, DMI series). Cultures were allowed to recover in the recording chamber for at least 2 minutes before recordings.

## Author Contributions

Larissa Breuer performed iAstrocyte differentiation, RNA extraction and initial analysis of RNAseq data, culturing, immunostaining and quantitation of iAstrocyte polyclonal and monoclonal cell-type specific markers and support of iNeurons, immunostaining and imaging of synapse markers in iNeuron, iNeuron/iAstro, and iNeuron/mAstro cultures, transduction of iAstrocytes with gfaABC1D-lck-GCaMP6f for calcium imaging and wrote the manuscript. Hanna Dubrovska plated and maintained iNeurons alone, iNeurons with iAstrocytes, and iNeurons with mouse astrocytes on HD-MEAs, performed weekly recordings, and analyzed recordings with Brainwave 5.6 software (3Brain), and performed RNAseq analysis and data visualization. Jeremy Krohn prepared mouse astrocytes and performed calcium imaging and analysis of iAstrocytes. John Carl Begley performed calcium imaging and analysis of iNeurons with iAstrocytes and iNeurons with mouse astrocytes. Hana Sheldon performed culturing, immunostaining and imaging of iNeuron/iAstro, and iNeuron/mAstro co-cultures. Katarzyna Ludwik designed iAstrocyte gene constructs, performed gene editing, introduced compound E as a neuronal differentiation promoting agent, and supervised the project. Harald Stachelscheid supervised the project. Camin Dean supervised the project and wrote the manuscript. All authors edited the manuscript.

## Acknowledgements

We thank the BIH Core Unit pluripotent Stem Cells and Organoids for support. Specifically, Valeria Fernandez Vallone for providing support with hIPSC line generation and validation and for editing the manuscript, Regina Jahn for technical assistance in performing PCRs, genotyping and clone picking, and Judit Küchler for technical assistance with pre-differentiation and banking of NGN2 NSCs. We thank Kai Guth, Corben Dean, and Nikita Cijsouw for assistance with plating and maintaining iNeuron, iNeuron/iAstrocyte, and iNeuron/mouse Astrocyte cultures. We thank the Advanced Medical BIOimaging Core Facility of the Charité-Universitätsmedizin Berlin (AMBIO) for support in acquisition of the imaging data.

